# The extent of gender and race/ethnicity imbalance in infectious disease dynamics research

**DOI:** 10.1101/2024.12.19.629420

**Authors:** Juliana C. Taube, Alexes Merritt, Shweta Bansal

**Author notes:** These authors contributed equally (Juliana C. Taube, Alexes Merritt).

## Abstract

Publication practices accumulate to affect credibility and career advancement. Understanding authorship and citation practices is critical to addressing inequities. While citation bias has been demonstrated in several fields, it remains uncharacterized in infectious disease dynamics (IDD), a quantitative, interdisciplinary domain highly visible during the COVID-19 pandemic. We analyze IDD articles and their bibliographies from 2000-2019 using machine-learning algorithms to infer the gender and race/ethnicity of each article’s lead and senior author. We examine authorship and citation patterns by gender and racial group across geographic scales, including characterizing the author composition of each article’s bibliography relative to the field. Our analysis reveals persistent gender and race imbalances in IDD research. Man-authored and White-authored publications dominate the field, with little progress in racial diversification of US and UK publications over the last two decades. Woman-authored articles have the most representative citation practices but are undercited, especially when women are senior authors. In the US and UK, most citations feature White lead and senior authors, even when citing articles have lead or senior authors of color. These findings underscore the urgent need for more inclusive IDD research practices. We discuss possible mechanisms and solutions to create opportunities for researchers from underrepresented groups.

## Introduction

Diverse teams produce better science [1, 2] yet numerous barriers prevent underrepresented individuals from entering, flourishing, and staying in scientific research (e.g., [3–5]). As it stands, White men generally dominate scientific fields: while women compose 51% of the United States (US) population, for example, they only make up 35% of the US STEM workforce [6]. The gender disparity in STEM fields follows a scissor-shaped curve: women slightly outnumber men at the undergraduate level, but in later career stages (Ph.D. and beyond), men dominate, widening the gap between genders at a particularly visible professional level [7]. There are similar disparities in racial and ethnic representation within STEM fields [6, 8]. Identifying these inequities within each academic field is necessary to address field-specific equity issues and, ultimately, to improve national and international scientific output and individual research experiences.

Publication and citation practices are two critical markers of scientific success that are particularly vulnerable to bias and could be contributing to these disparities. These metrics are integral to visibility, credibility, career advancement, and attaining leadership positions but are not always objectively constructed and may be amplifying existing inequities [9]. Characterizing how publication rates and citation practices vary within the scientific ecosystem is one way to evaluate the diversity and equity within a scientific field. This work has been conducted across disciplines [10–12] and in several specific fields including political science [13], international relations [14,15], neuroscience [16–18], physics [19], astronomy [20], and medicine [21]. For example, Bertelero and colleagues’ found that White neuroscience authors preferentially cite other White authors and that neuroscience authors of color are being increasingly undercited despite authoring an increasing proportion of articles [17]. Likewise, Caplar and colleagues’ found that women are lead-authoring an increasing proportion of astronomy publications, but their work is undercited compared to men [20]. Within infectious disease research, a recent study on editors and authorship in 40 infectious disease journals found that women are underrepresented in senior author and editor positions and suggested that recruiting more women editors could help increase the publication rate for women [22]. The field of infectious disease dynamics is distinct from most infectious disease research due to its highly quantitative nature and is highly visible to the public due to its ability to provide actionable insights during health crises. These distinctions make the field both prone to gender and race disparities and a priority for ensuring equitable research methods. However, there have been no comprehensive analyses of publication and citation practices in infectious disease dynamics to date.

Here, we address this gap by examining and quantifying the author composition and citation practices by gender and race/ethnicity within the interdisciplinary and impactful field of infectious disease dynamics. We compile a dataset of over 10,000 articles in infectious disease dynamics and infer the gender and race/ethnicity of lead (first) and senior (last) authors of these articles using validated machine-learning packages. We test three hypotheses: (1) men and White (lead/senior) authors are over-represented in articles in the field (especially in the Global North); (2) even after accounting for the composition of the field, articles with women (lead/senior) authors are undercited compared to articles authored by men authors; and (3) articles by women and authors of color exhibit more representative citation practices than articles with men and White (lead/senior) authors. Our findings characterize the degree of inequity in infectious disease dynamics publication and citation practices and suggest potential ways to remediate these issues.

## Methods

### Data collection

Infectious disease dynamics (IDD) is an interdisciplinary field that publishes in a broad range of disciplinary and general scientific journals. Therefore, it is challenging to define the boundaries of the published literature in IDD by selecting journals or conducting keyword searches. Similarly, because no IDD international professional organization exists, membership in the field is not feasible to identify. For our analysis, we thus identified authors in the IDD field by identifying the articles that cite a set of influential primary research and review IDD articles before 2020. To identify this set of influential research, we searched the Web of Science Core Collection (WoSCC) and selected articles that developed or discussed dynamical models of population-scale disease transmission and had accrued an average of 50 or more citations per year since publication. This resulted in 23 research articles published between 1990 and 2015. (Additional details and the the complete list of articles can be found in the Supplement.) To characterize authorship patterns in IDD, we define the *authorship dataset* as the set of articles published in 2000-2019 in WoSCC that cite these influential articles as of November 25, 2024 and are cited by another authorship dataset article by November 25, 2024.

We also define an alternative definition for the IDD authorship dataset by identifying articles that have cited three seminal IDD books: *Infectious Diseases of Humans: Dynamics and Control* [23], *Modeling Infectious Diseases in Humans and Animals* [24], and *Mathematical Tools for Understanding Infectious Disease Dynamics* [25]. These articles were extracted on April 13, 2023 from WoSCC. We present our findings based on this alternative definition in the Supplement (Figure S1).

To characterize citation patterns in IDD, we define the *citation dataset* as the set of articles cited in the articles of the authorship dataset. To exclude irrelevant citations, citation dataset articles must either be in the authorship dataset or have been cited sufficiently based on their date of publication (from more than 2 times for 2019, to more than 5 times for 2000, determined using the 90th percentile of the number of citations for other articles published the same year). We perform sensitivity analyses on this threshold for the citation dataset (Figures S19, S20).

### Inferring identities

We inferred the gender and race/ethnicity of the lead and senior authors of each article in the authorship and citation datasets using the machine-learning algorithms implemented in genderize.io [26,27] and ethnicolr [28], respectively. Genderize.io (https://genderize.io/) is designed to infer the gender of a first name using a proprietary algorithm. The response is either male, female, or none, as well as a confidence probability, representing the number of data entries used to calculate the response and the proportion of names with the gender returned in the response. The underlying data (of over a billion individuals) is collected from social networks across 79 countries and 89 languages. Ethnicolr (https://github.com/appeler/ethnicolr) is a Python package designed to predict race and ethnicity based on the sequence of characters in a person’s name. The package uses character-level recurrent neural networks (RNNs) to make these predictions using either first and last name or just last name. The model is trained on multiple data sources, including US Census data (2000 and 2010), Florida voting registration data, and Wikipedia data. The predictions are broad race/ethnicity categories based on US definitions (White, Black, Asian, Hispanic), as well as a probability score for each potential category. Only identities classified with over a 70% confidence were included; we performed sensitivity analyses on this threshold (Figures S15, S16, S17, S18). We also assessed the sensitivity of our results to the choice of algorithm by additionally using namsor [29] for classification (Figures S13, S14). The accuracy of all three algorithms has been validated and these algorithms are regularly used in academic research [30–36]. We additionally use the affiliation for the senior author to assign a home country to each article, allowing us to specify our analysis to certain countries (e.g., US, UK) or the Global North (which is broadly comprised of Northern America, Europe, Israel, Japan, South Korea, Australia, and New Zealand). All geographically restricted analyses exclude articles that are missing senior author country affiliations. For articles with missing author first names or non-delineated first and last names, we pulled additional author name information from the CrossRef API, if available. To disambiguate authors who are recognized by different names across articles or with only initials for first names (i.e., E. Halloran vs Elizabeth Halloran), we used the algorithm in [16] to identify authors in the dataset with first and last names matching the initials and last names, and replaced initials with the inferred full name. If there were different full author names that match a set of initials, then we did not replace the name (i.e., Elizabeth Halloran and Edward Halloran).

### Analyses

To understand authorship patterns, we measured the proportion of articles in the authorship dataset authored by each gender and race/ethnicity group across time and geography. When analyzing mentorship patterns we removed any articles with only a single author. We performed bootstrapping (1000 draws) to address small sample sizes. These proportions are compared relative to the gender and race/ethnicity representation of the general population and STEM workforce, where possible, in the respective geography (e.g., global, Global North, US, UK).

To characterize gendered citation practices, we calculated the cumulative fraction of articles of each lead and senior authorship gender type (man-man (MM), woman-woman (WW), woman-man (WM), man-woman (MW)) across time; this distribution represents the pool of articles available to cite in a given bibliography. We then compared the makeup of each bibliography with that of the field one month prior to publication to assess which author types were being over- or undercited. We included citations of articles written before 2000 by assuming that the composition of the field had not changed before 2000. To understand whether the identity of the citing authors affected their citation practices, we also calculated the proportion of each gender pairing in each bibliography for articles with men lead and senior authors versus articles with a woman lead and/or senior author. We bootstrapped both of these estimates by randomly selecting articles from the authorship dataset 1000 times and analyzing their bibliographies. We repeat the same analyses for race/ethnicity with a focus on White-White and lead or senior person of color (POC)-authored articles due to low sample size of mixed race authorship. We restricted this analysis to years with 25 or more citations. In all our analyses of citation practices, we exclude self-citations to measure broader scholarly impact, reduce sub-disciplinary biases [37], and prevent inflation of diversity biases [38]. In our supplementary analyses, we consider the sensitivity of our findings to this decision (Figures S21, S22).

We further analyzed citation practices using a linear regression model to understand how lead and senior author identity predict citation rate (defined as the ratio of the number of total citations of an article and the number of years since publication). We exclude self-citations and control for 2023 Journal Impact Factor (JIF) and whether an article was published by authors in the Global North. (See Supplement for sensitivity analyses and diagnostics; articles missing senior author affiliation country are excluded.) The model was structured as:

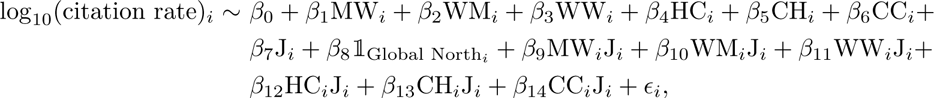

where M stands for man, W for woman, C for person of color, H for White, and J for the square root of 2023 Journal Impact Factor, and *ɛ_i_*∼ *N* (0*, σ*^2^).

## Results

We identify 10,660 articles in the authorship dataset and 16,195 unique articles for the citation dataset cited in 220,570 citations across the bibliographies of the authorship dataset. The published literature of the IDD field grew rapidly and consistently across the last two decades (Figure 1A) driven primarily by articles lead- and senior-authored by men and White individuals (in the US and UK) (Figures S6, S7, S9).The authorship dataset spans 101 countries of affiliation, with the majority of articles (65%) published between 2000 and 2019 written by senior authors from the Global North (Figure 1B). The two countries with the most published articles per population size were the UK with 938 articles (14 per million people) and the US with 2,855 articles (8.70 per million people) (Figure S2). We were able to infer senior author gender for 85% of authorship articles and 86% of unique citation articles and senior author race for 63% of authorship articles and 53% of unique citation articles.

**Figure 1.**
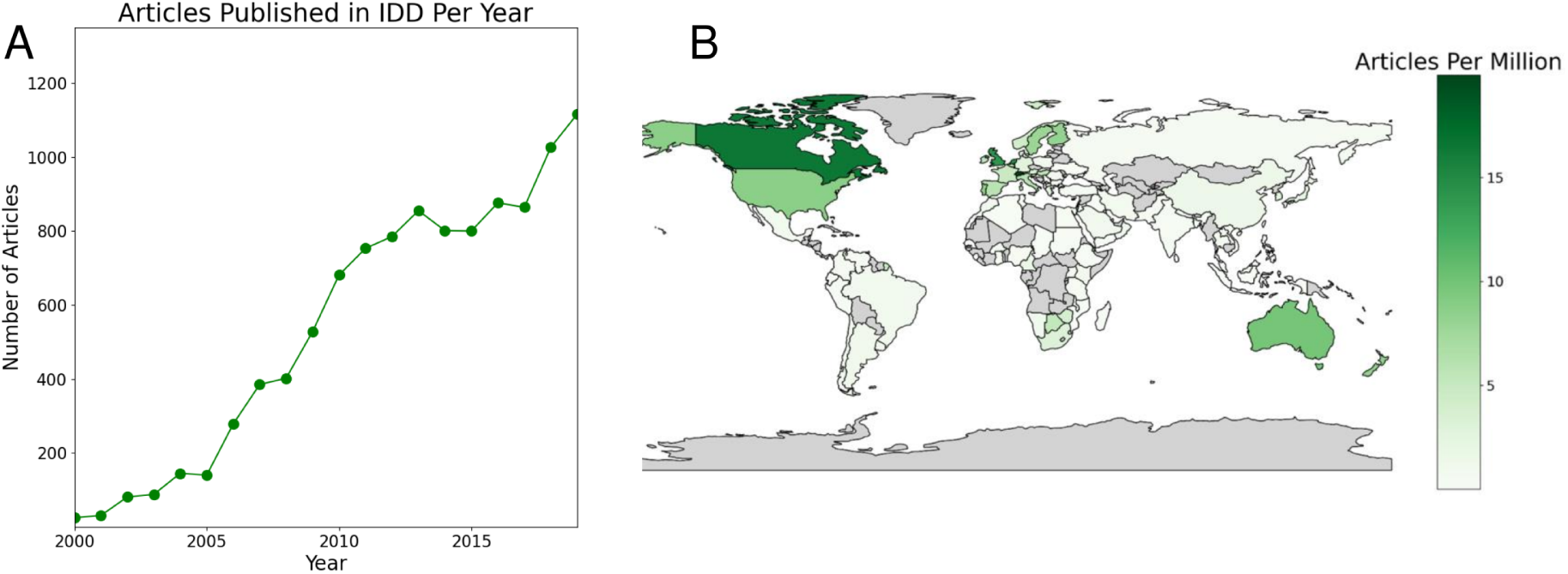
Infectious disease dynamics (IDD) publications have rapidly increased in the last two decades, but most senior authors are concentrated in the Global North. (A) Number of infectious disease dynamics publications per year from 2000 to 2019. (B) Distribution of articles published per million (general) population by country from 2000 to 2019, based on the affiliations of the senior authors.

### IDD publication authorship is dominated by men while woman-authored articles are undercited

We find that men lead-authored 61% of articles and senior-authored 67% of the articles published in IDD from 2000 to 2019, despite men composing only 50.4% of the global population. The ratio of man-authored to woman-authored articles has remained relatively constant since the early 2000s for both lead and senior authors (Figures 2A, S5). Concerning mentorship, 43% of articles have men lead and senior authors while only 6% of articles have women lead and senior authors. These ratios are heavily skewed compared to the global population and equal gender representation within the field, but do not indicate unequal mentoring practices after taking into account the field’s disproportionate number of men authors (Figure 2B). These findings are consistent across geographic scales, gender-inference approaches, field definitions, and after accounting for uncertainty in our sample (Figures S10, S11, S13, S3).

**Figure 2.**
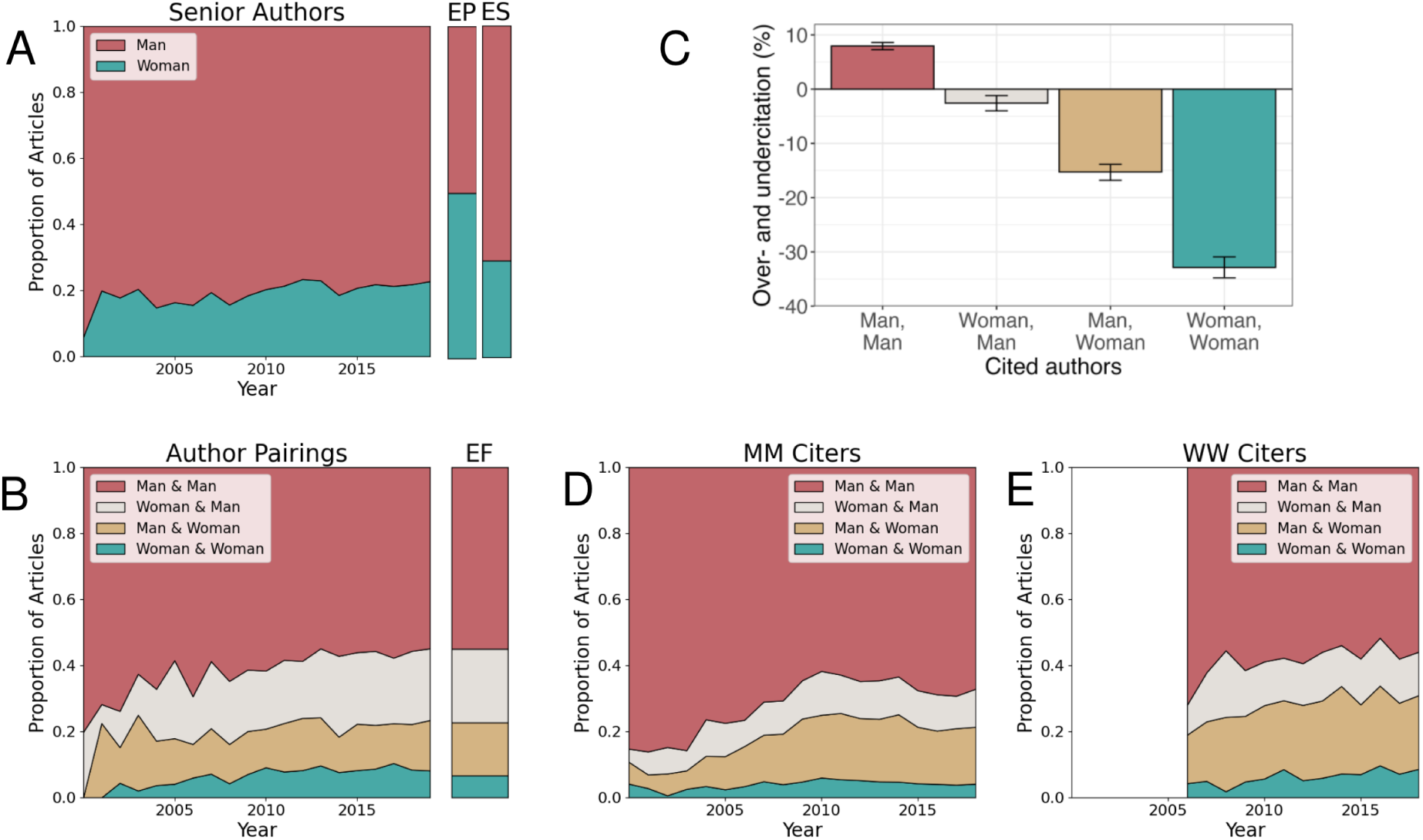
Globally, infectious disease dynamics publications are predominantly authored by men, but mentorship practices across genders do not appear biased. Women are undercited, particularly by man-man authored articles. (A) Proportion of articles senior-authored by each gender from 2000 to 2019. Expected proportions based on (global) population (EP) in 2019 and based on the Global STEM workforce (ES) in 2023 [39] are shown in the color bars at right. (B) Proportion of articles authored by a given (lead & senior author) gender pair from 2000 to 2019. Expected proportions based on the field (EF) in 2019 are shown in the color bar at right; these expectations are based on the observed proportions in Figure S5. (C) Rates of over- and undercitation by article lead and senior author gender based on authorship gender composition prior to publication date. (D) Authorship gender breakdown of articles cited in articles by men lead and senior authors (MM) from 2000 to 2019. (E) Authorship gender breakdown of articles cited in articles written by women lead and senior authors (WW) from 2006 to 2019.

In the context of citation practices, women lead- and senior-authored articles are undercited and the woman’s authorship position affects the degree of undercitation. WM articles are undercited by 3%, MW by 15%, and WW by 33% (Figure 2C). These findings are robust to geography and gender inference package (Figures S10, S13) but are sensitive to the definition of the IDD field (Figure S11); the rate of undercitation of WW authored articles is not different from zero in the alternative definition of the field that has a smaller sample size. In 2019, 66% of references in MM articles are also MM-authored publications and this trend has been fairly consistent over time (Figure 2D). WW articles have had increasingly gender-diverse bibliographies with only 55% of their citations corresponding to MM articles in 2019 (Figure 2E). MM articles consistently cite articles with women as lead and/or senior authors at a lower rate than WW articles cite articles with women as lead and/or senior authors (statistically significant for each year with a t-test). However, MM articles are increasingly citing articles with women as lead and/or senior authors at a slightly higher rate compared to the rate at which WW articles cite articles with women lead and/or senior authors (slope=0.011, intercept=0.21 for MM citers and slope=0.006, intercept=0.360 for WW citers with a Mann-Kendall test with p-value < 0.05). In Figure S21, we test the sensitivity of these findings to the inclusion of self-citations and find the results to be qualitatively robust.

### Race/ethnicity disparities are present in authorship and citation practices

Because race is a social construct that is experienced and treated differently across contexts, we focus on the race/ethnicity of authors in the Global North, where race is understood similarly. We restrict our analysis to four race/ethnicity categories in the US and UK (White, Black, Hispanic, and Asian) so that we can compare authorship composition with the general population and STEM workforce of each country. We find that most US publications have White authors with growing Asian authorship, which is less diverse than the general population (Figure 3A, S8). Similarly, nearly all UK publications have White authors, but authorship more closely reflects the general population (Figure 3B, S8). These findings are consistent across definitions of the IDD field, race/ethnicity inference methods, and after accounting for uncertainty in our sample (Figures S12, S14, S4).

**Figure 3.**
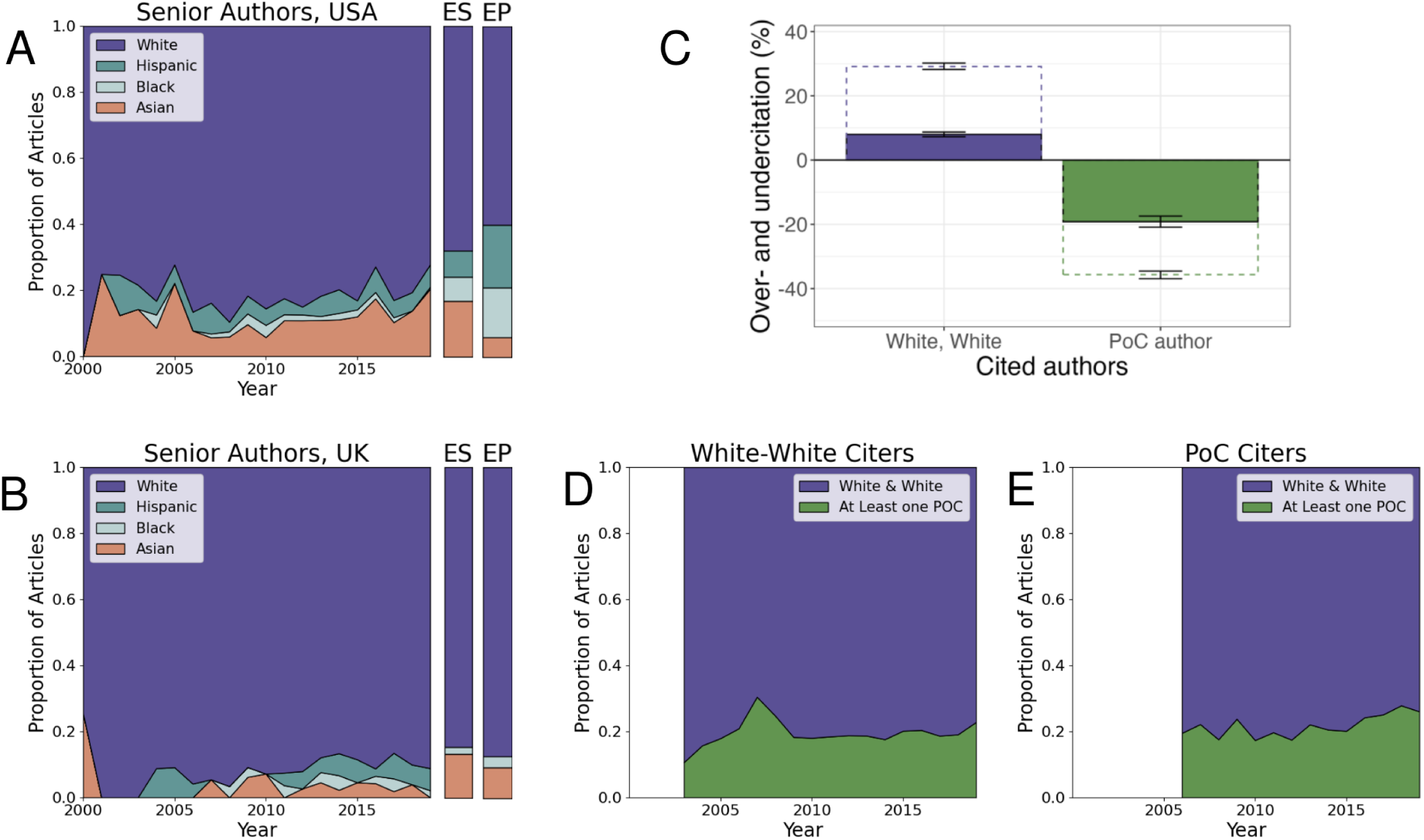
Authorship in infectious disease dynamics is not racially diverse and White-authored articles are overcited compared to articles that have a non-White lead and/or senior author. (A) Proportion of articles senior-authored by each race/ethnicity group in the USA from 2000 to 2019. Expected values (ES) based on the national STEM workforce based on 2019 National Science Foundation reporting [40], and expected values (EP) based on 2019 Census data [41] are shown in the color bars to the right. (B) Proportion of articles senior-authored by each race/ethnicity group in the UK from 2000 to 2019. Expected values (ES) based on the national STEM academic staff [42] in 2019, expected values (EP) based on 2019 Office of National Statistics Data [43] are shown in the color bars to the right. (C) Rates of over- and undercitation by article lead and senior author race/ethnicity based on authorship race/ethnicity composition prior to publication date. PoC author denotes that the lead and/or senior author was a person of color; sample size was too small to disaggregate by author position for PoC authors. Filled boxes denote results for articles published by authors with affiliations within the Global North citing other work from the Global North. Dashed boxes denote results for articles published by authors with affiliations within the Global North citing articles with any affiliation location. (D) Racial/ethnic composition of articles cited in articles written by White lead and senior authors in the Global North, citing other articles authored in the Global North from 2003 to 2019. (E) Racial/ethnic composition of articles cited in articles written by a lead and/or senior author of color in the Global North, citing other articles authored in the Global North from 2006 to 2019.

Likewise, citation practices show significant biases. Articles with a lead and/or senior author of color are undercited, while articles with White lead and senior authors are overcited in the Global North (Figure 3C). The rates of over- and undercitation are more extreme when author’s bibliographies are expected to reflect global race/ethnicity authorship (dashed boxes in Figure 3C). Articles authored by lead and/or senior PoC generally cite articles authored by lead and/or senior PoC at a higher rate than articles with White lead and senior authors cite lead and/or senior PoC authored articles (8 of 14 years show statistical significance with a t-test, p-value <0.005). Additionally, both groups of authors are increasingly citing articles with authors of color as lead and/or senior authors at similar rates (White-White citers: slope=0.0031, intercept=0.15; PoC citers: slope=0.0060, intercept=0.15 with a Mann-Kendall test with p-value < 0.05) (Figure 3D, E). In Figure S22, we test the sensitivity of these findings to the inclusion of self-citations and find the results to be qualitatively robust.

### Author race & gender influence citation rate after controlling for publication venue

We found that author gender pairing and race pairing are significantly associated with article citation rate, even after accounting for the impact factor of the journal where the article was published (Figure 4). Woman-woman, PoC-White, and PoC-PoC articles had significantly lower citation rates, while Journal Impact Factor significantly increased citation rate. However, woman-woman, PoC-White, and PoC-PoC publications in high-impact factor journals had higher citation rates relative to man-man and White-White publications, though these interactions were not statistically significant. Sensitivity analyses varying geography, self-citation inclusion, and citation thresholds for inclusion can be found in the Supplement (Figures S25, S26, S27, S28, S29, S30, S31, S32).

**Figure 4.**
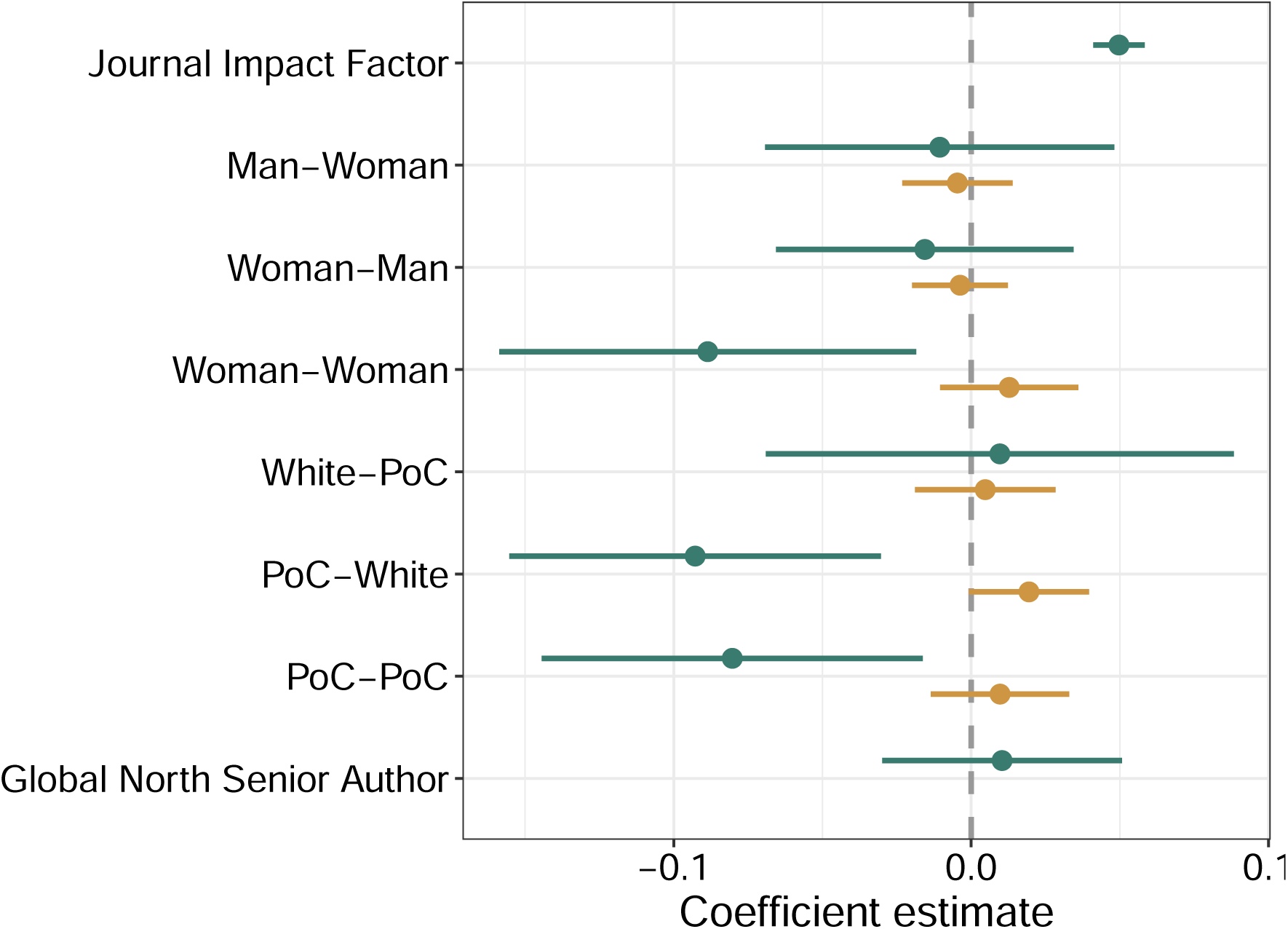
Citation rates in infectious disease dynamics are gender- and race/ethnicity-biased. Coefficient estimates are shown from a linear model predicting citation rate (num. citations/years since publication) based on article and author information. Teal point-ranges show the main effects (coefficient estimate and 95% confidence interval) for the independent impact of the predictor on citation rates, and gold point-ranges show the interaction effects for the impact of the predictor on citation rates for different values of Journal Impact Factor. Coefficients represent the difference from citation of man-man and White-White authored articles with non-Global North senior authors and average Journal Impact Factor. Woman-woman, PoC-White, and PoC-PoC authored articles are significantly less cited, while articles in high-impact journals are significantly more cited. Journal impact factor is square-root transformed to ensure normality. PoC stands for person of color.

## Discussion

Infectious disease dynamics is a rapidly growing area of interdisciplinary research. As the field reflects on lessons learned from the early stages of the COVID-19 pandemic and identifies areas of future work, it is paramount to consider which voices are underrepresented and underrecognized in the field. This introspection is essential for promoting inclusivity, but also because publications and citations are key ways that academic success, credibility, and potential are measured, and public dissemination is achieved. We address this pressing issue by comprehensively analyzing the authorship and bibliographic landscape of infectious disease dynamics publications. We overcome challenges to define the field by curating a list of influential articles and analyze the relevant articles citing these influential works. We use validated machine-learning packages [44] to infer the gender and race/ethnicity identity of the authors of each article and conduct sensitivity analyses on this identity inference, our definition of the field, and using varying spatial scales to demonstrate that our results are robust.

We find that publication and citation practices in the field of infectious disease dynamics are notably biased. Most publications are authored by men or White authors which doesn’t reflect the composition of the general population (see intersectional results in Figure S33). Work authored by men or White scientists is overcited even after accounting for their disproportionate composition of the field. In contrast, work authored by women or people of color is heavily undercited, and this amplifies with gender or race assortativity (i.e., mentorship by authors of the same identities). Author position also affects the rate of undercitation, with greater bias when women are in more senior authorship positions. The same trend may apply for authors of color, but we did not have sufficient sample size to detect it. Our regression analysis shows that these issues persist even after accounting for Journal Impact Factor. However, we do learn that citation rates of the articles by women or people of color in high-impact journals are higher relative to man-man and White-White authored articles, respectively. We speculate that this might reflect the phenomenon of overperformance for underrecognized individuals, in which systemically disadvantaged individuals are driven to and held to a higher standard for reaching the top of their field [45].

These gender results are consistent across spatial scales, but our race/ethnicity findings show greater geographic variability. The United States and the United Kingdom produced the most articles in the Global North, but authorship within these countries is not representative of the racial/ethnic makeup of the general population. Citation practices by race/ethnicity are similarly biased. Articles authored by White lead and senior authors are overcited while articles with at least one author of color are undercited by more than 30%. This difference cannot be explained by White lead and senior authors citing fewer authors of color (Figure 3D,E).

We recognize that our interpretation of gender as binary and race/ethnicity fitting into four classifications is limiting and prevents analysis of some of the most vulnerable scientists, including, but not limited to, non-binary individuals and Indigenous populations. Furthermore, race/ethnicity categorization can meaningfully differ across countries. Using names to infer gender and race can be inaccurate and ignores physical forms of identity expression [46]. However, names and affiliations are the key identifying information that editors, reviewers, and authors have when choosing to read, accept, review, or cite a given article. Thus, inaccuracies in gender or race/ethnicity inference may well reflect how an unknown author is perceived by others. We focus on authors listed first and last, ignoring shared authorship positions and the identity makeup of the full authorship list. We acknowledge that our sample may be missing some IDD publications or include non-IDD work, but we must start to address these issues of inequity even in the absence of perfect data. Despite these limitations, our findings align with those from other disciplines, such as neuroscience, physics, astronomy, and medicine [16,19–21]. In particular, similar to the fields of mathematics and computer science, we find an increasing percentage of articles authored by women, but not at a sufficient rate to equal the rate at which men are authoring articles [47,48]. Likewise, our findings are nearly identical to those in software engineering and computer science, more broadly, where man-authored articles are overcited and woman-authored articles are undercited, with variation by woman author position [49, 50]. However, woman-woman authored articles in software engineering and computer science overcite other woman-woman authored articles to a greater degree than in our study. Analyses of citation practices by race/ethnicity in quantitative fields are limited, though across published US authors of any field, Freeman and Huang observed assortativity in collaborations based on race/ethnicity [51]. Thus, the field of infectious disease dynamics appears to have demographic composition and citation practices similar to those of other quantitative disciplines.

As the field of infectious disease dynamics moves forward with additional urgency and scrutiny due to the COVID-19 pandemic, it is imperative to take action to increase the diversity and equity in the field [52]. We must also recognize how the pandemic has likely only exacerbated many of the disparities we have documented here [53]. For example, women had lower research productivity than men [54], including submitting fewer manuscripts during the early months of the pandemic [55]. We advocate for increased journal editorial board and reviewer diversity [22, 56–58] to leverage the effect of homophily. Additionally bibliographic transparency [19], in which editors and authors ensure that the composition of submitted bibliographies reflects the gender and race/ethnicity composition of the field, is critical. To facilitate this process, we’ve created a tool so that authors can check their own bibliographies relative to expected compositions for the field of IDD [59]. Additionally, individuals should consider citing both older initial work and recent applications by more junior and diverse authors in their manuscripts, a practice that can be aided by the removal of reference length limits by journals. Finally, academia should consider broadening the definitions of scientific contributions to reflect the variety of work that advances the field but may not be traditionally valued, for example, by acknowledging the value of dashboard, app, and software package creation, science communication, and policy engagement. To move forward, the field must assess and examine our current composition and practices so that we can measure future progress; we begin such an effort here and call on others to join us in acknowledging and correcting the inequities within our field.

## Acknowledgments

We appreciate data sharing and valuable technical support provided by Ann Beynon and Rob Pritchett of Clarivate. We are grateful to Simon Frost for providing statistical expertise and assistance. We also thank Eva Rest and Andrew Tiu for their technical assistance and Juliet Pulliam, Anne Cori, Amy Wesolowski for their valuable feedback on this work.

## Data availability

This project utilizes a commercial dataset provided by Clarivate and the raw data cannot be republished. For further information, please contact Clarivate. The data and code used in our analysis is provided for reproducibility at https://github.com/bansallab/IDD_imbalance.

## Supplementary Information

## Appendix A: Defining the field

To define the authorship dataset, we identify a set of influential articles in the field of infectious disease dynamics. To do this, we searched the Web of Science Core Collection for articles matching the search terms in Table S1. This resulted in 69,654 articles. We then excluded all articles that had an average annual citation rate below 50 citations per year, which resulted in 275 articles. We then excluded articles that were not relevant to the field (excluded journal names and title terms can be found in Table S1). The resulting list has 23 articles published from 1990 to 2015 and can be found in Table S2.

**Table S1.**
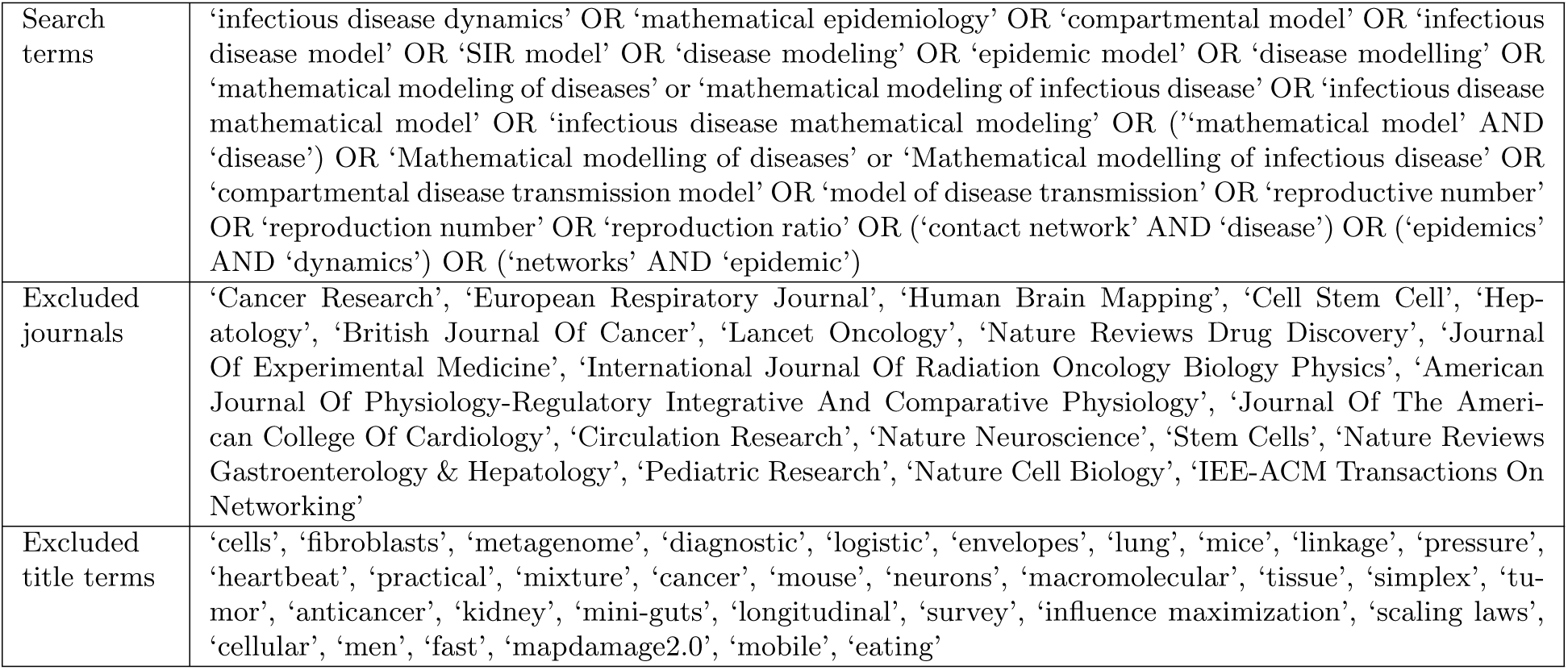
Web of Science Core Collection search terms and exclusion criteria to identify set of 23 influential articles in infectious disease dynamics.

|  |  |
| --- | --- |
| Search terms | 'infectious disease dynamics' OR 'mathematical epidemiology' OR 'compartmental model' OR 'infectious disease model' OR 'SIR model' OR 'disease modeling' OR 'epidemic model' OR 'disease modelling' OR 'mathematical modeling of diseases' or 'mathematical modeling of infectious disease' OR 'infectious disease mathematical model' OR 'infectious disease mathematical modeling' OR ('mathematical model' AND 'disease') OR 'Mathematical modelling of diseases' or 'Mathematical modelling of infectious disease' OR 'compartmental disease transmission model' OR 'model of disease transmission' OR 'reproductive number' OR 'reproduction number' OR 'reproduction ratio' OR ('contact network' AND 'disease') OR ('epidemics' AND 'dynamics') OR ('networks' AND 'epidemic') |
| Excluded journals | 'Cancer Research', 'European Respiratory Journal', 'Human Brain Mapping', 'Cell Stem Cell', 'Hepatology', 'British Journal Of Cancer', 'Lancet Oncology', 'Nature Reviews Drug Discovery', 'Journal Of Experimental Medicine', 'International Journal Of Radiation Oncology Biology Physics', 'American Journal Of Physiology-Regulatory Integrative And Comparative Physiology', 'Journal Of The American College Of Cardiology', 'Circulation Research', 'Nature Neuroscience', 'Stem Cells', 'Nature Reviews Gastroenterology & Hepatology', 'Pediatric Research', 'Nature Cell Biology', 'IEEE-ACM Transactions On Networking' |
| Excluded title terms | 'cells', 'fibroblasts', 'metagenome', 'diagnostic', 'logistic', 'envelopes', 'lung', 'mice', 'linkage', 'pressure', 'heartbeat', 'practical', 'mixture', 'cancer', 'mouse', 'neurons', 'macromolecular', 'tissue', 'simplex', 'tumor', 'anticancer', 'kidney', 'mini-guts', 'longitudinal', 'survey', 'influence maximization', 'scaling laws', 'cellular', 'men', 'fast', 'mapdamage2.0', 'mobile', 'eating' |

**Table S2.** List of influential seed papers for defining the authorship dataset.

| Authors | Article Title | Journal | Publication Year | Citation Rate |
| --- | --- | --- | --- | --- |
| Diekmann, O. et al | On The Definition And The Computation Of The Basic Reproduction Ratio $R_0$ In Models For Infectious-Diseases In Heterogeneous Populations | Journal Of Mathematical Biology | 1990 | 97.50 |
| Hethcote, H.W. et al | The Mathematics Of Infectious Diseases | Siam Review | 2000 | 171.17 |
| Pastor-Satorras, R. et al | Epidemic Spreading In Scale-Free Networks | Physical Review Letters | 2001 | 182.04 |
| Van Den Driessche, P. et al | Reproduction Numbers And Sub-Threshold Endemic Equilibria For Compartmental Models Of Disease Transmission | Mathematical Biosciences | 2002 | 276.59 |
| Newman, M.E.J. et al | Spread Of Epidemic Disease On Networks | Physical Review E | 2002 | 112.45 |
| Eubank, S. et al | Modelling Disease Outbreaks In Realistic Urban Social Networks | Nature | 2004 | 67.05 |
| Castillo-Chavez, C. et al | Dynamical Models Of Tuberculosis And Their Applications | Mathematical Biosciences And Engineering | 2004 | 59.40 |
| Lloyd-Smith, J.O. et al | Superspreading And The Effect Of Individual Variation On Disease Emergence | Nature | 2005 | 87.21 |
| Ferguson, N.M. et al | Strategies For Containing An Emerging Influenza Pandemic In Southeast Asia | Nature | 2005 | 70.32 |
| Keeling, M.J. et al | Networks And Epidemic Models | Journal Of The Royal Society Interface | 2005 | 63.47 |
| Ferguson, N.M. et al | Strategies For Mitigating An Influenza Pandemic | Nature | 2006 | 84.28 |
| Altizer, S. et al | Seasonality And The Dynamics Of Infectious Diseases | Ecology Letters | 2006 | 60.33 |
| Wallinga, J. et al | How Generation Intervals Shape The Relationship Between Growth Rates And Reproductive Numbers | Proceedings Of The Royal Society B-Biological Sciences | 2007 | 57.00 |
| Chitnis, N. et al | Determining Important Parameters In The Spread Of Malaria Through The Sensitivity Analysis Of A Mathematical Model | Bulletin Of Mathematical Biology | 2008 | 57.75 |
| Granich, R.M. et al | Universal Voluntary Hiv Testing With Immediate Antiretroviral Therapy As A Strategy For Elimination Of Hiv Transmission: A Mathematical Model | Lancet | 2009 | 98.33 |
| Fraser, C. et al | Pandemic Potential Of A Strain Of Influenza A (H1N1): Early Findings | Science | 2009 | 95.40 |
| Balcan, D. et al | Multiscale Mobility Networks And The Spatial Spreading Of Infectious Diseases | Proceedings Of The National Academy Of Sciences Of The United States Of America | 2009 | 59.73 |
| Diekmann, O. et al | The Construction Of Next-Generation Matrices For Compartmental Epidemic Models | Journal Of The Royal Society Interface | 2010 | 78.14 |
| Funk, S. et al | Modelling The Influence Of Human Behaviour On The Spread Of Infectious Diseases: A Review | Journal Of The Royal Society Interface | 2010 | 58.21 |
| Brockmann, D. et al | The Hidden Geometry Of Complex, Network-Driven Contagion Phenomena | Science | 2013 | 76.55 |
| Granell, C. et al | Dynamical Interplay Between Awareness And Epidemic Spreading In Multiplex Networks | Physical Review Letters | 2013 | 67.91 |
| Cori, A. et al | A New Framework And Software To Estimate Time-Varying Reproduction Numbers During Epidemics | American Journal Of Epidemiology | 2013 | 56.91 |
| Pastor-Satorras, R. et al | Epidemic Processes In Complex Networks | Reviews Of Modern Physics | 2015 | 265.11 |

**Figure S1.**
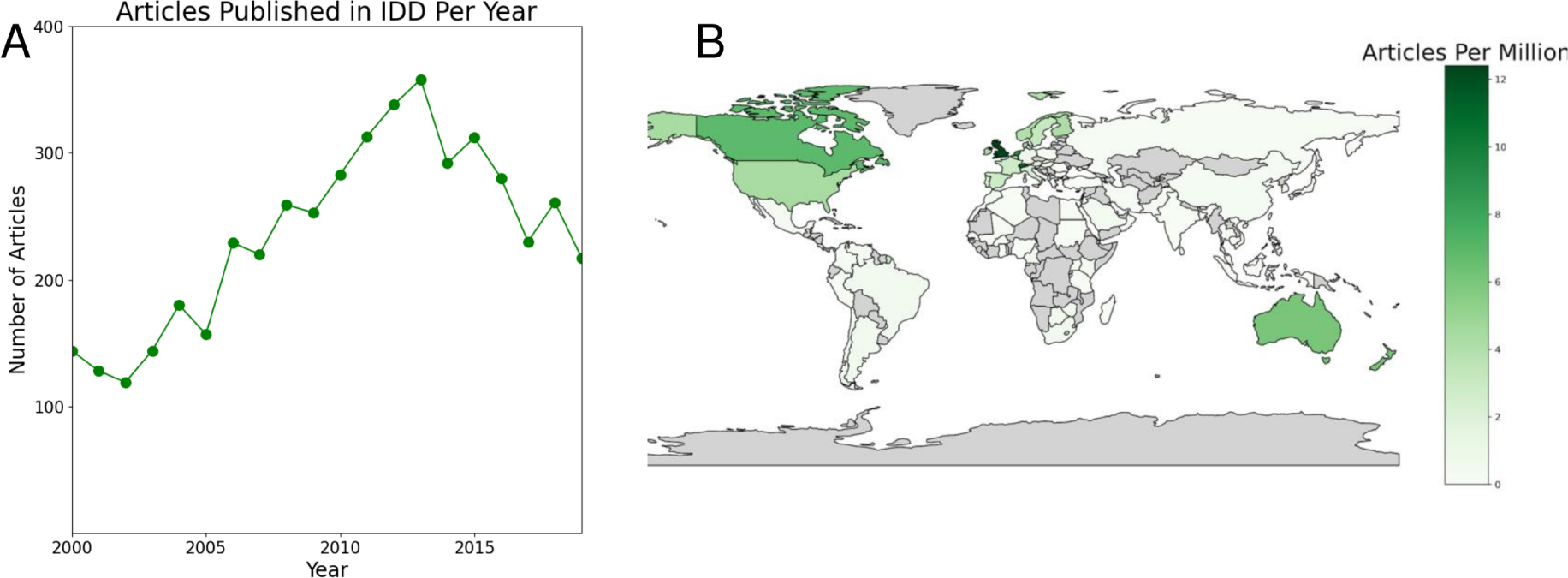
Number of articles by year based on alternative definition of the field. Here, the field is defined as articles citing [23–25]. (A) Number of articles published per year globally. (B) Number of articles published per million residents in 2019.

**Figure S2.**
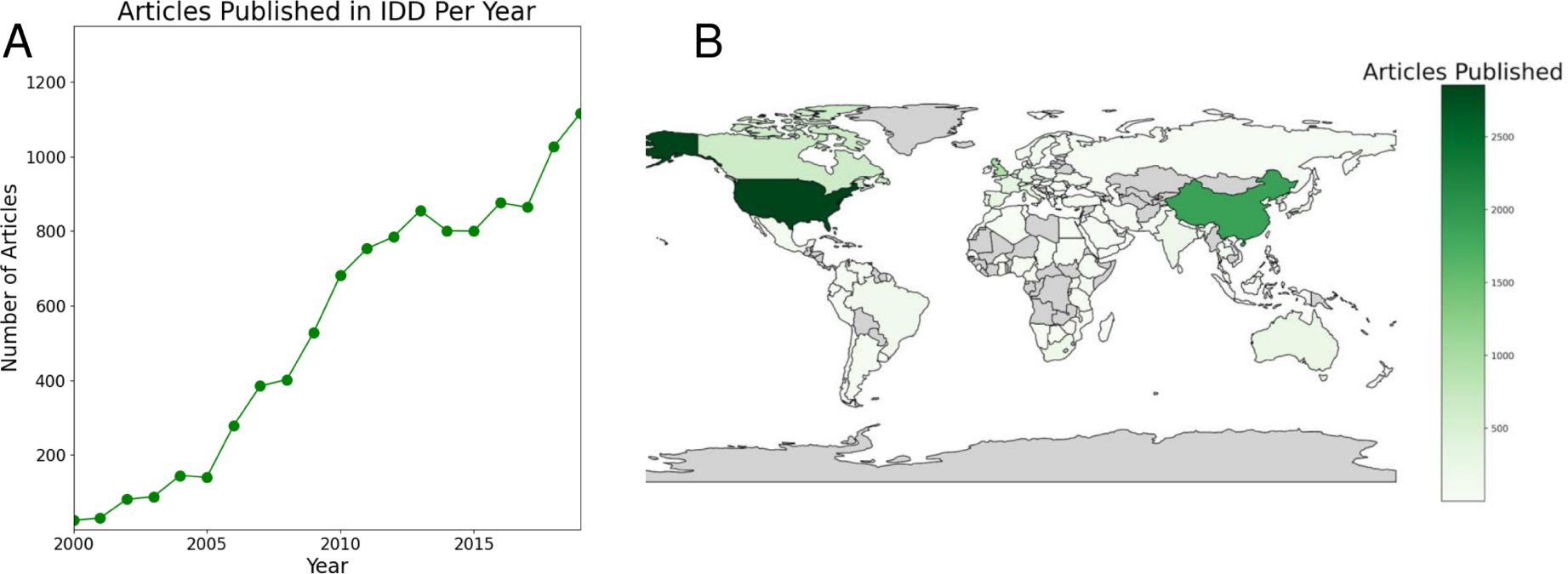
Total number of articles published in the field of infectious disease dynamics (IDD) from 2000-2019.

## Appendix B: Uncertainty in authorship and citation practices

**Figure S3.**
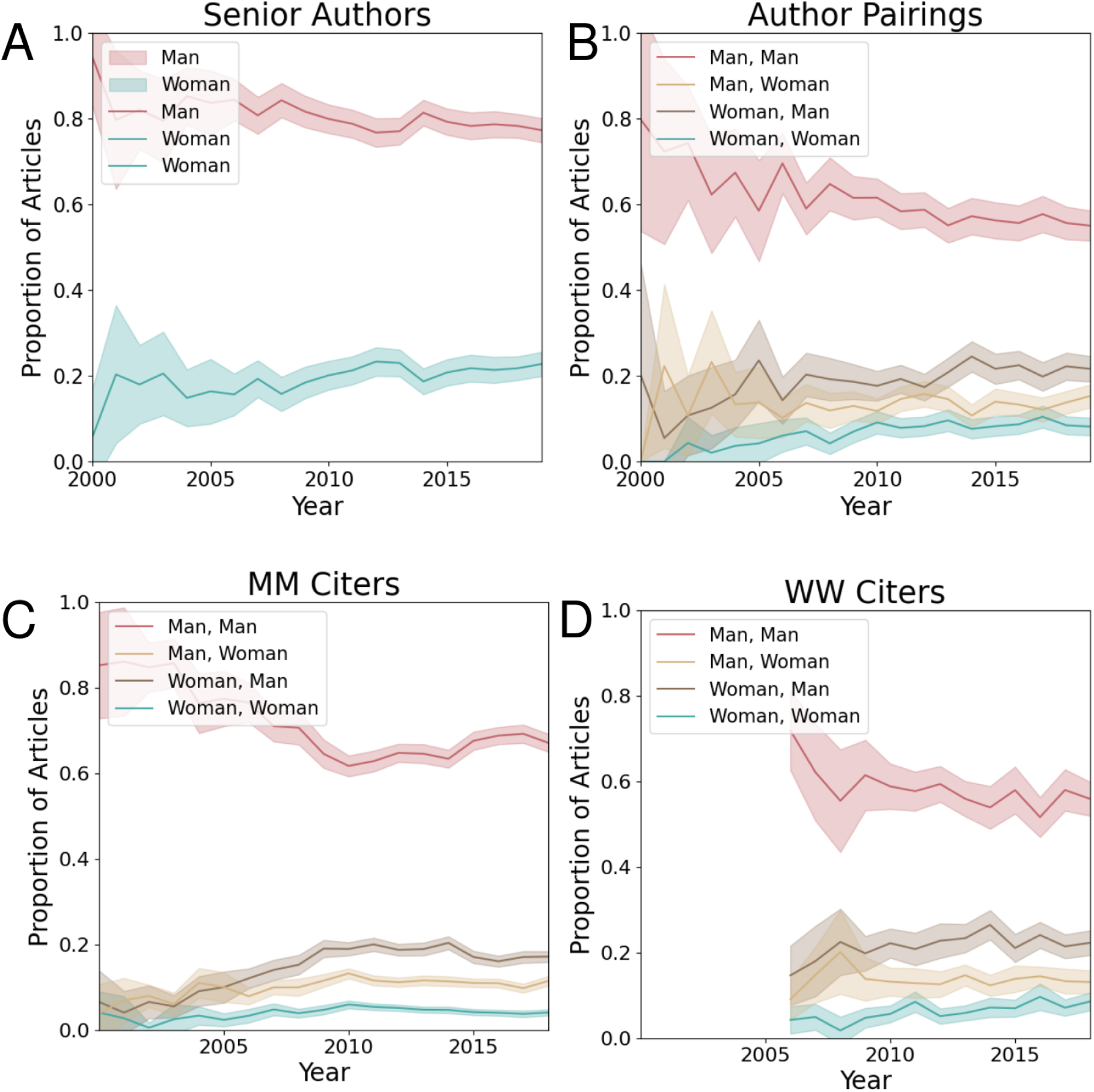
Gendered authorship and citation practices of the field with 2 standard deviations from the mean. (A) Proportion of articles senior-authored by each gender from 2000 to 2019, with shaded region representing 2 standard deviations from the mean. (B) Proportion of articles authored by a given (lead & senior author) gender pair from 2000 to 2019, with shaded region representing 2 standard deviations from the mean. (C) Authorship gender breakdown of articles cited by men lead and senior authors (MM), with shaded region representing 2 standard deviations from the mean from 2000 to 2019. (D) Authorship gender breakdown of articles cited by women lead and senior authors (WW), with shaded region representing 2 standard deviations from the mean from 2006 to 2019.

**Figure S4.**
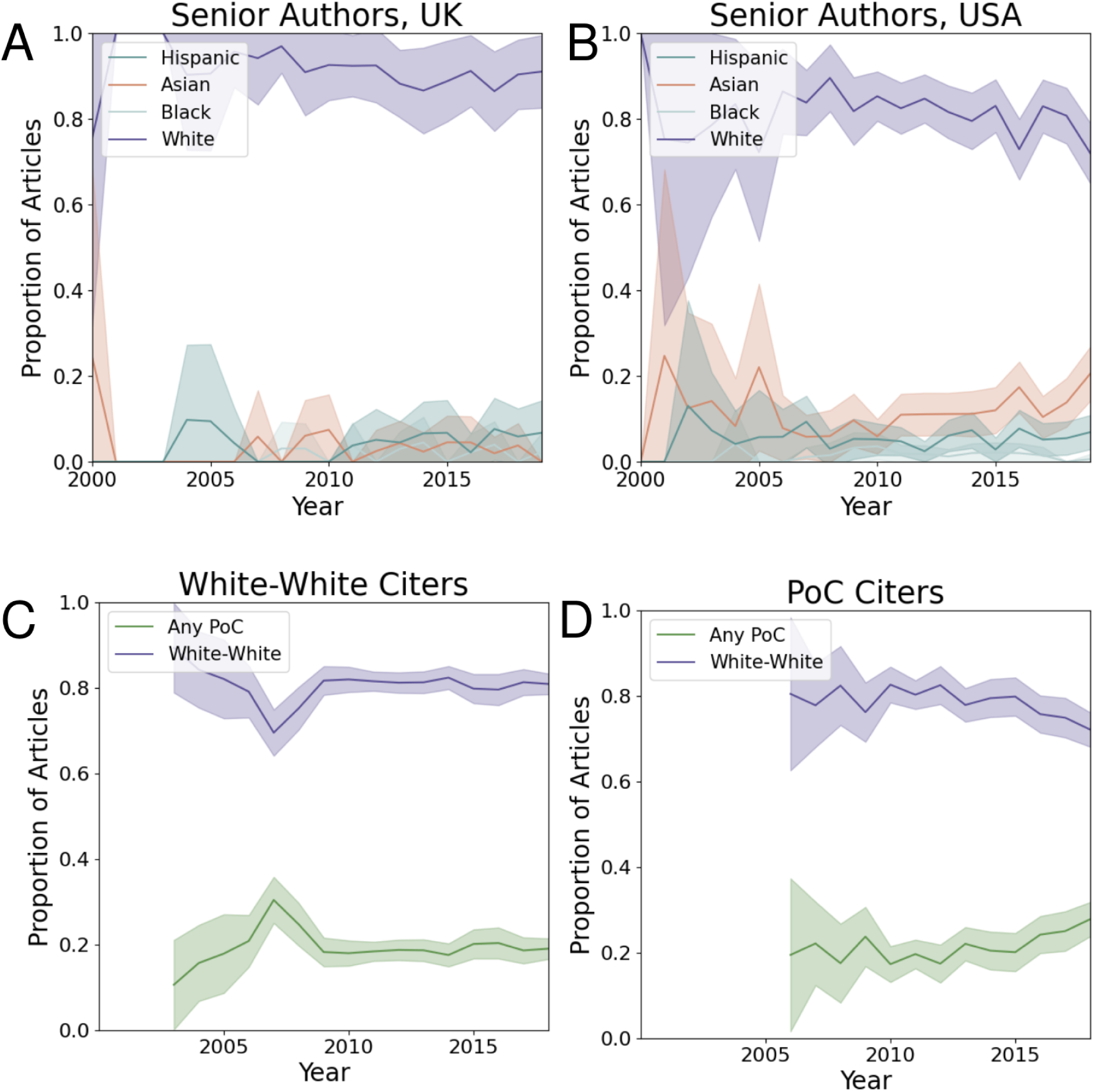
Race/ethnicity authorship and citation practices of the field with 2 standard deviations from the mean. (A) Proportion of articles senior-authored by each race/ethnicity group in the USA from 2000 to 2019, with shaded region representing 2 standard deviations from the mean. (B) Proportion of articles senior-authored by each race/ethnicity group in the UK from 2000 to 2019, with shaded region representing 2 standard deviations from the mean. (C) Racial/ethnic composition of articles cited by White lead and senior authors in the Global North, citing other articles authored in the Global North, with shaded region representing 2 standard deviations from the mean. (D) Racial/ethnic composition of articles by a lead and/or senior author of color in the Global North, citing other articles authored in the Global North, with shaded region representing 2 standard deviations from the mean.

## Appendix C: Demographic comparison of lead to senior author

**Figure S5.**
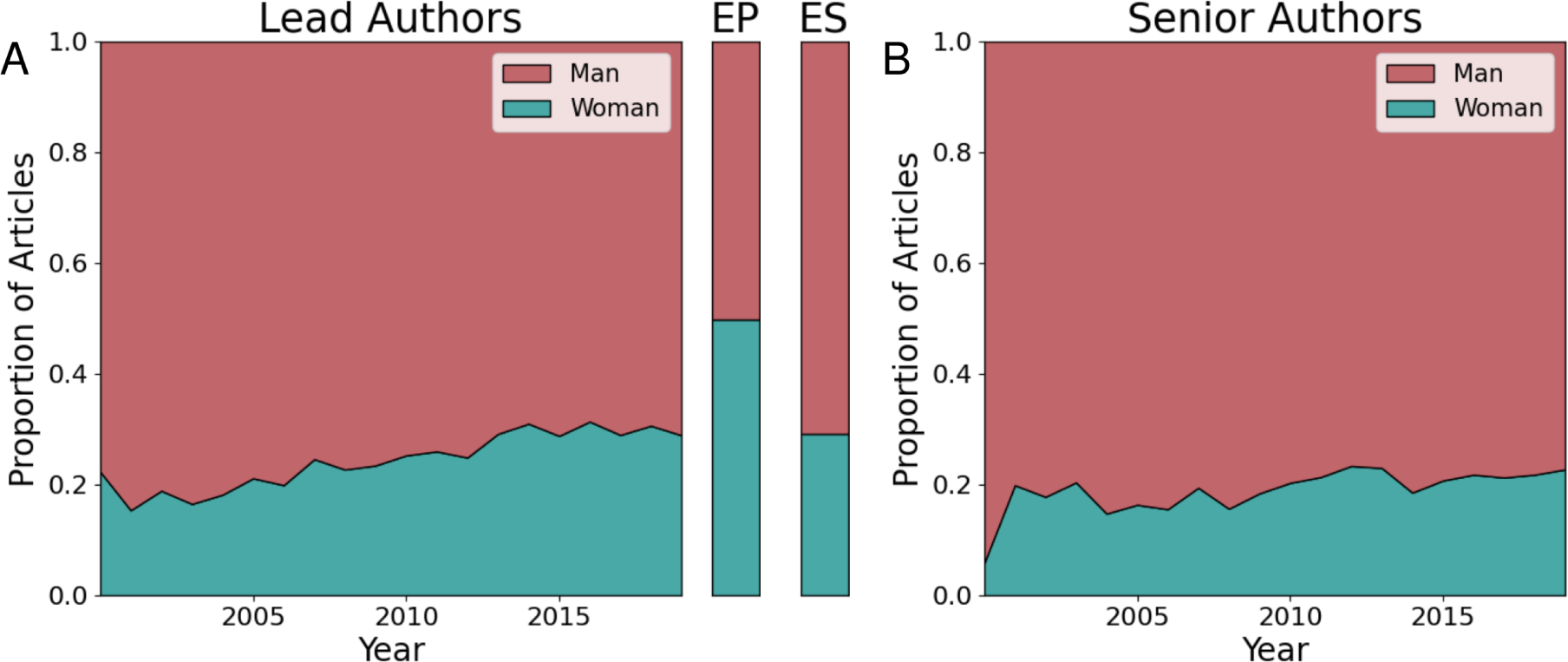
Globally, lead authors are more representative of gender demographics than senior authors. (A) Proportion of articles lead-authored by each gender from 2000 to 2019. Expected values (EP) for 2019 based on the global population are shown in the color bar at right. (B) Proportion of articles senior-authored by each gender from 2000 to 2019.

**Figure S6.**
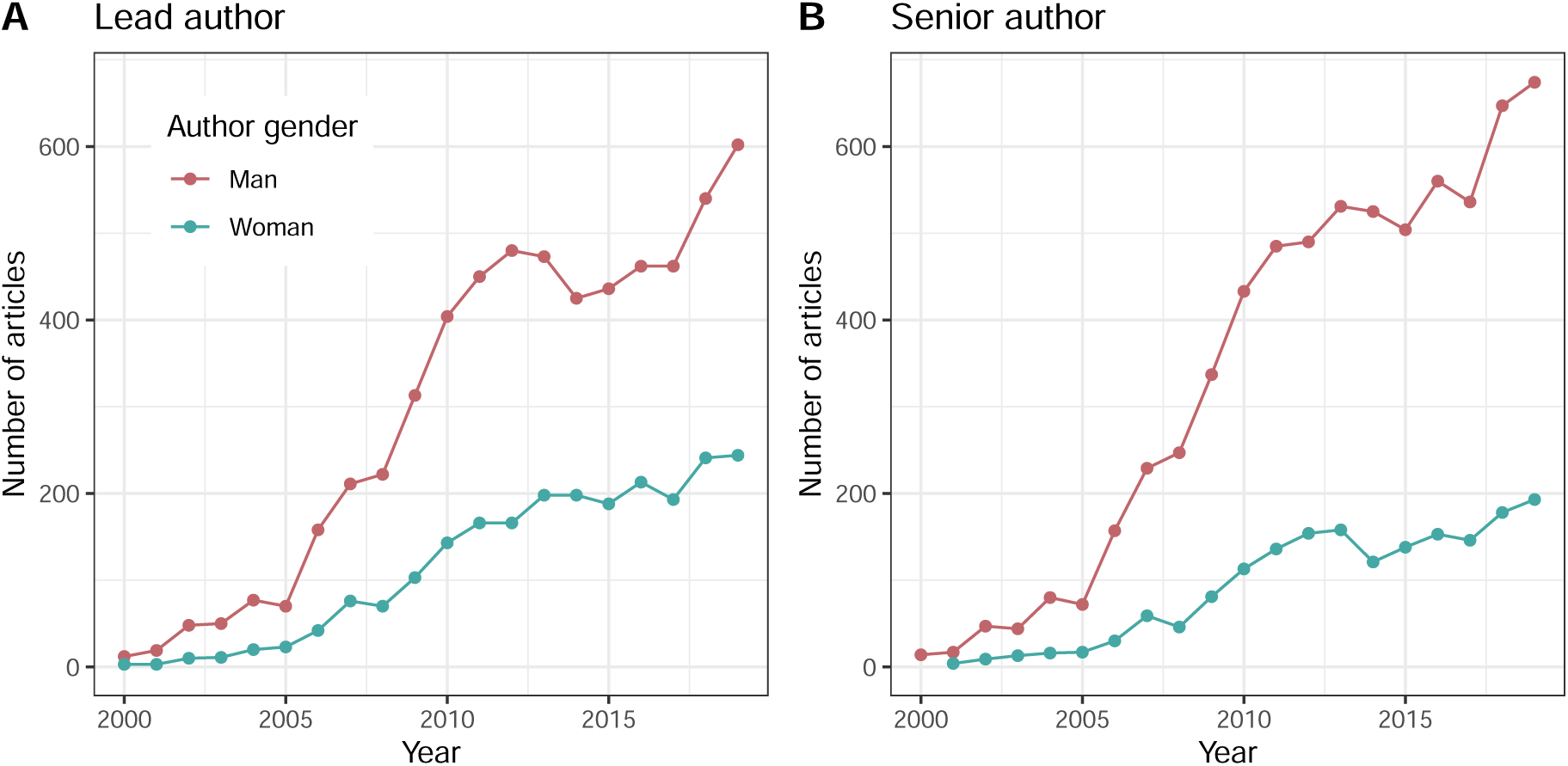
Articles with men lead or senior authors are growing faster than articles with woman lead or senior authors. Number of articles each year (A) lead- or (B) senior-authored by each gender.

**Figure S7.**
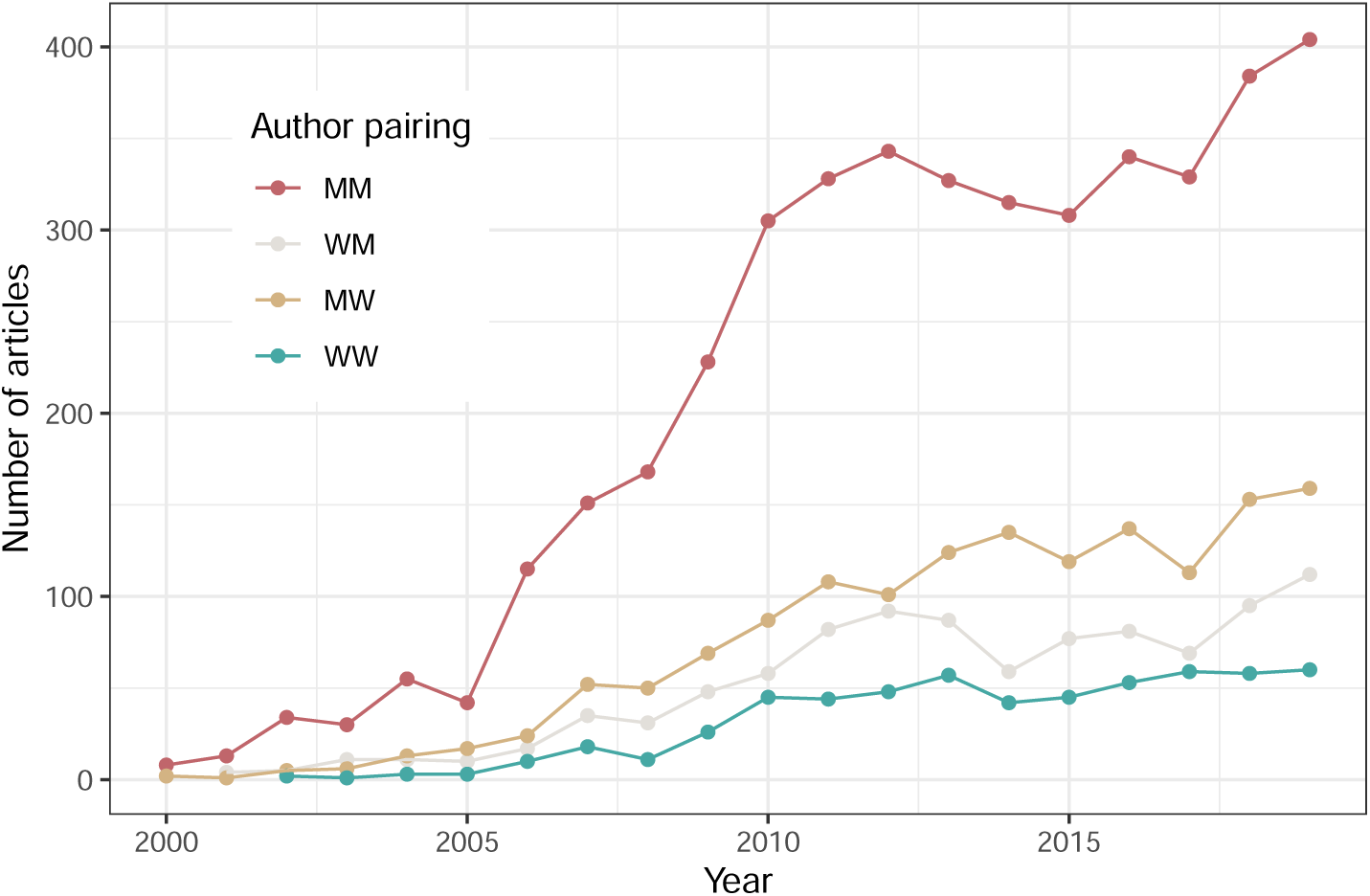
Increase in articles over time is driven by articles with men as lead and senior authors. Number of articles each year by lead and senior author gender.

**Figure S8.**
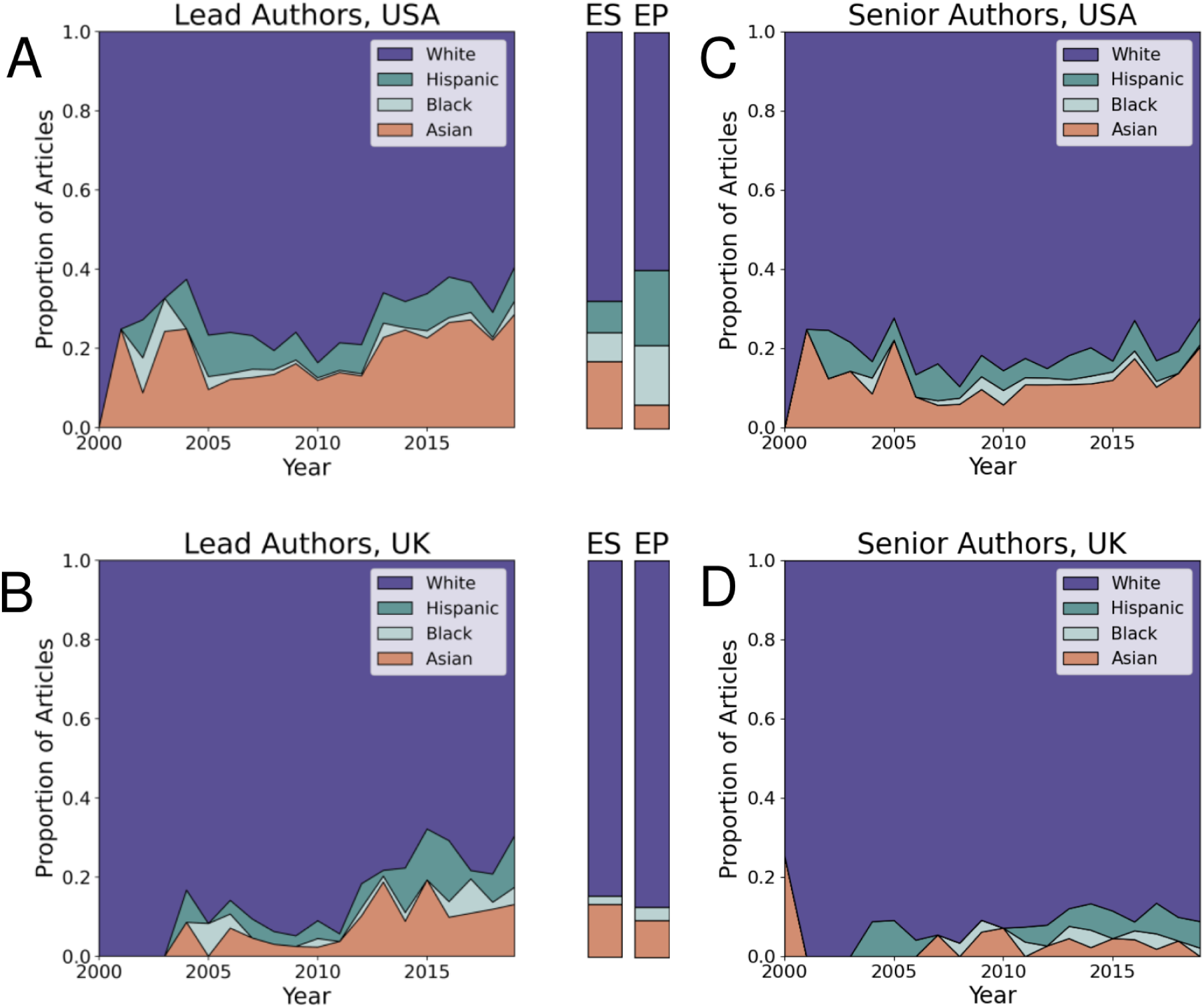
In both the USA and UK, lead authors are more diverse than their senior counterparts. (A) Proportion of articles lead-authored by each race/ethnicity group in the USA from 2000 to 2019. Expected values (ES) based on the national STEM workforce based on 2019 National Science Foundation reporting [40], and expected values (EP) based on 2019 Census data [41] are shown in the color bars in the middle. (B) Proportion of articles lead-authored by each race/ethnicity group in the UK from 2000 to 2019. Expected values (ES) based on the national STEM academic staff [42] in 2019, expected values (EP) based on 2019 Office of National Statistics Data [43] are shown in the color bars to the right. (C) Proportion of articles senior-authored by each race/ethnicity group in the USA from 2000 to 2019. (D) Proportion of articles senior-authored by each race/ethnicity group in the UK from 2000 to 2019.

**Figure S9.**
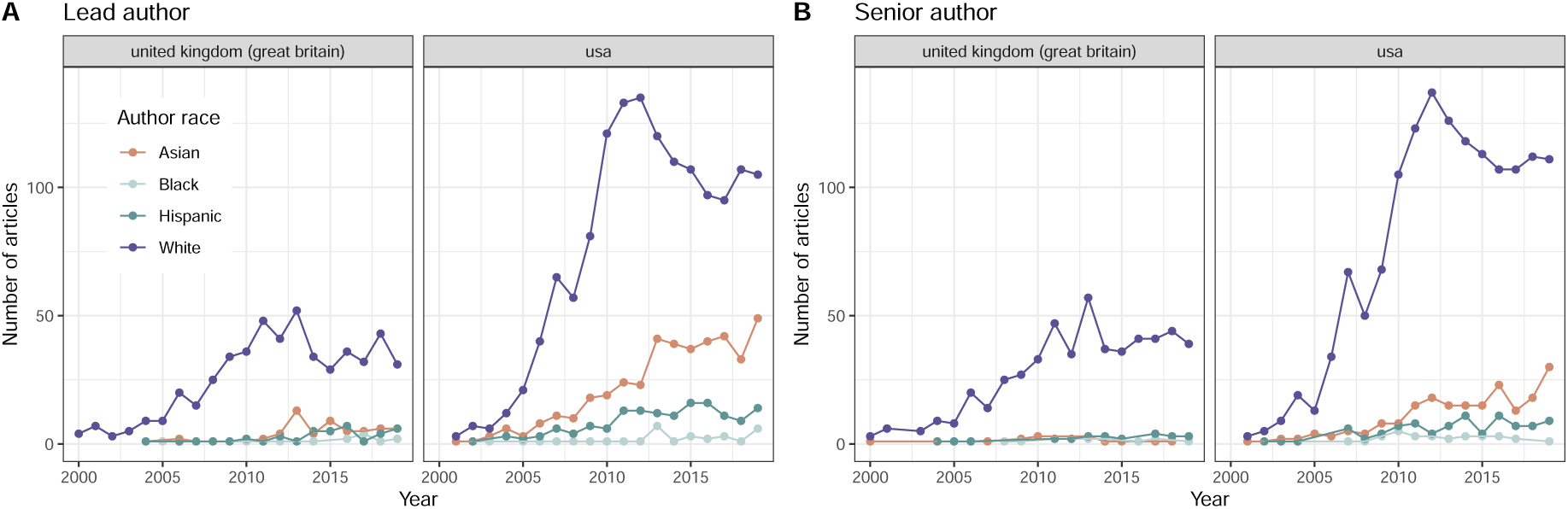
Number of articles each year authored by each race/ethnicity by author position and country. (A) Lead author race/ethnicity for the UK and US. (B) Senior author race/ethnicity for the UK and US.

## Appendix D: Gender results for the Global North

**Figure S10.**
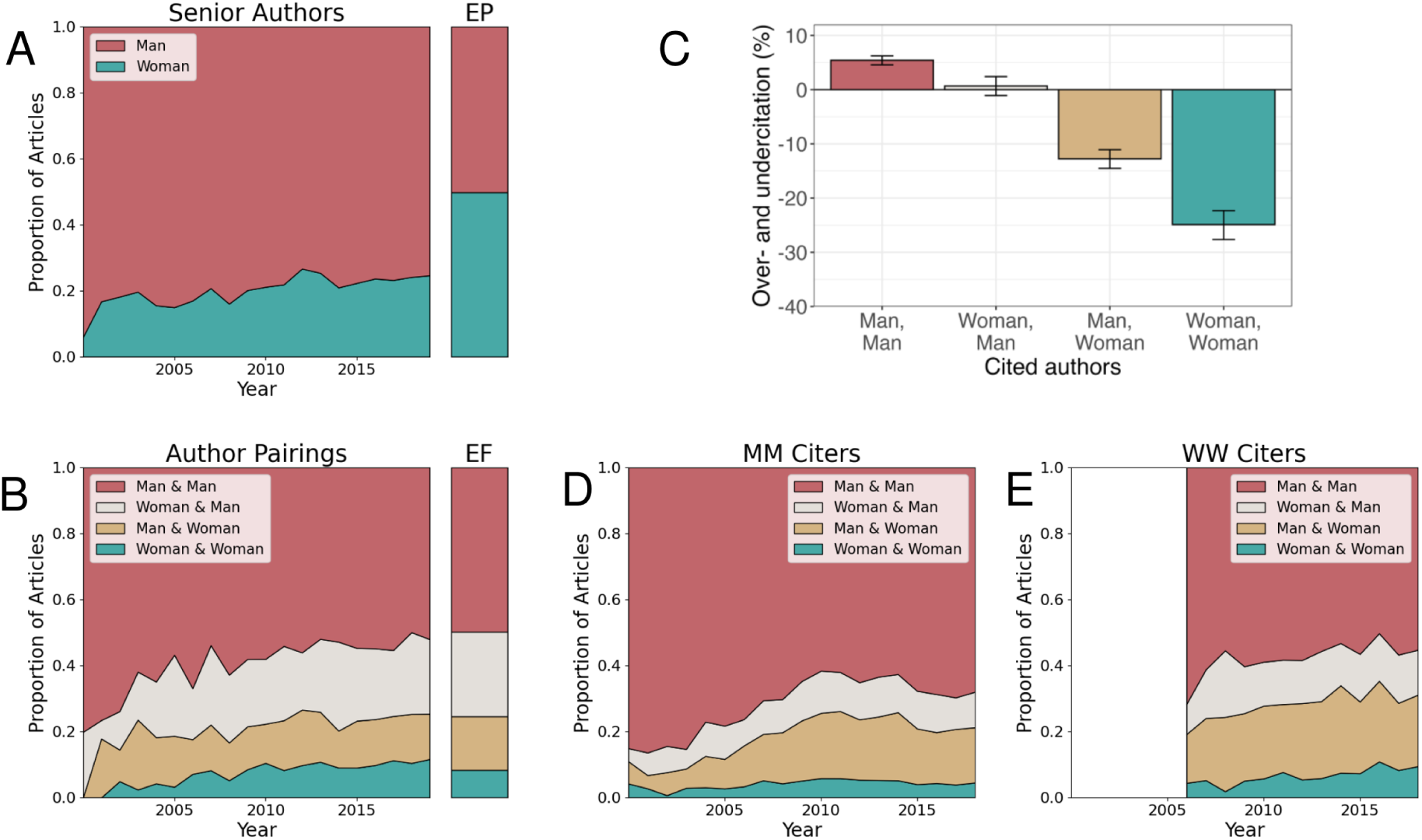
Authorship and citation practices in the Global North mirror global patterns with respect to gender. (A) Proportion of articles senior-authored by each gender from 2000 to 2019. Expected proportions based on (Global North) population (EP) in 2019 and based on the Global STEM workforce (ES) in 2023 [39] are shown in the color bars at right. (B) Proportion of articles authored by a given (lead & senior author) gender pair from 2000 to 2019. Expected proportions based on the field (EF) in 2019 are shown in the color bar at right. (C) Rates of over- and undercitation by article lead and senior author gender based on authorship gender composition prior to publication date. (D) Authorship gender breakdown of articles cited in articles by men lead and senior authors from 2000 to 2019.(E) Authorship gender breakdown of articles cited in articles written by women lead and senior authors from 2006 to 2019.

## Appendix E: Results based on different definition of the field

**Figure S11.**
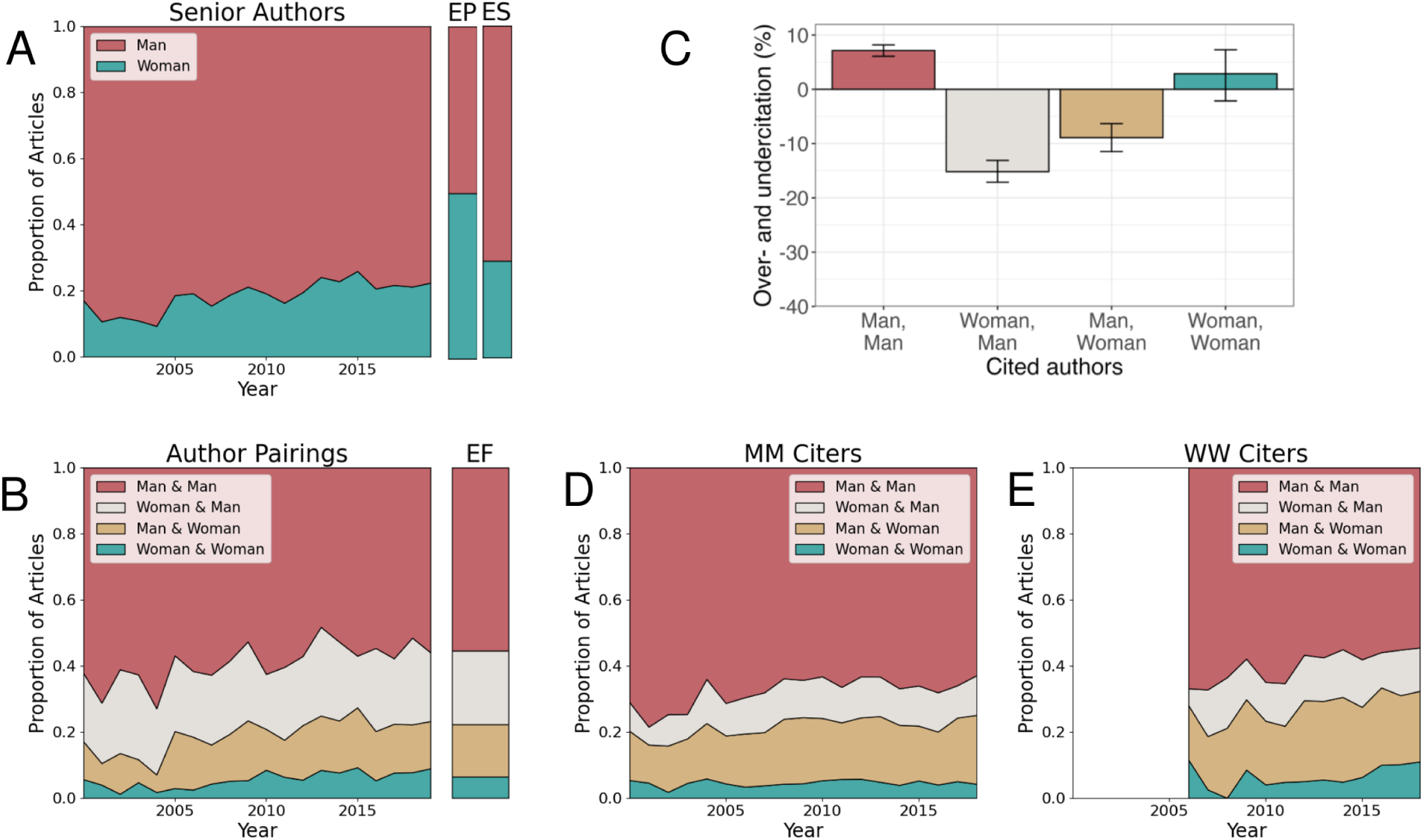
Authorship patterns remain largely unchanged, with some deviations in citation patterns as a result of changes in definition of the field. Here, the field is defined as articles citing [23–25]. (A) Proportion of articles senior-authored by each gender from 2000 to 2019. Expected values (EP) based on global proportions in 2019 and based on the Global STEM workforce (ES) in 2023 [39] are shown in the color bars at right. (B) Proportion of articles authored by a given (lead & senior author) gender pair from 2000 to 2019. Expected values (EF) produced based on the gender composition of the field in 2019 are shown in the color bar at right.(C) Rates of over- and undercitation by article lead and senior author gender based on authorship gender composition prior to publication date. (D) Authorship gender breakdown of articles cited in articles by men lead and senior authors from 2000 to 2019.(E) Authorship gender breakdown of articles cited in articles written by women lead and senior authors from 2006 to 2019.

**Figure S12.**
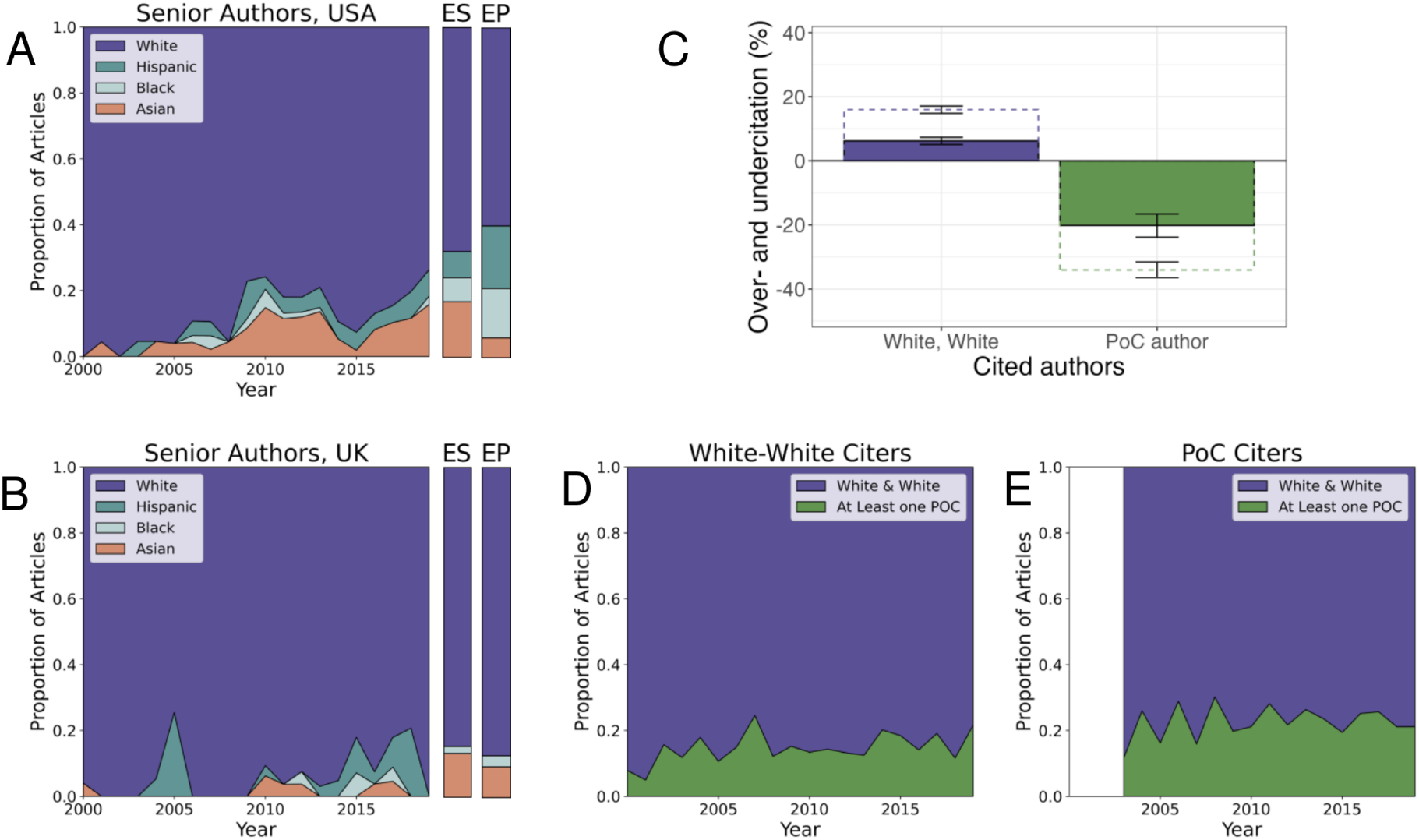
Authorship and citation practices remained biased towards White authors even with a different definition of the field. Here, the field is defined as articles citing [23–25]. (A) Proportion of articles senior-authored by each race/ethnicity group in the USA from 2000 to 2019. Expected values (ES) based on the national STEM workforce based on 2019 National Science Foundation reporting [40], and expected values (EP) based on 2019 Census data [41] are shown in the color bars to the right. (B) Proportion of articles senior-authored by each race/ethnicity group in the UK from 2000 to 2019. Expected values (ES) based on the national STEM academic staff [42] in 2019 and expected values (EP) based on 2019 Office of National Statistics Data [43] are shown in the color bars to the right. (C) Rates of over- and undercitation by article lead and senior author race/ethnicity based on authorship race/ethnicity composition prior to publication date. PoC author denotes that the lead and/or senior author was a person of color; sample size was too small to disaggregate by the author of color’s position. Filled boxes denote results for articles published by authors with affiliations within the Global North citing other work from the Global North. Dotted boxes denote results for articles published by authors with affiliations within the Global North citing articles with any affiliation location. (D) Racial/ethnic composition of articles cited in articles written by White lead and senior authors in the Global North, citing other articles authored in the Global North from 2000 to 2019. (E) Racial/ethnic composition of articles cited in articles written by a lead and/or senior author of color in the Global North, citing other articles authored in the Global North from 2003 to 2019.

## Appendix F: Results based on different gender/race inference tools

**Figure S13.**
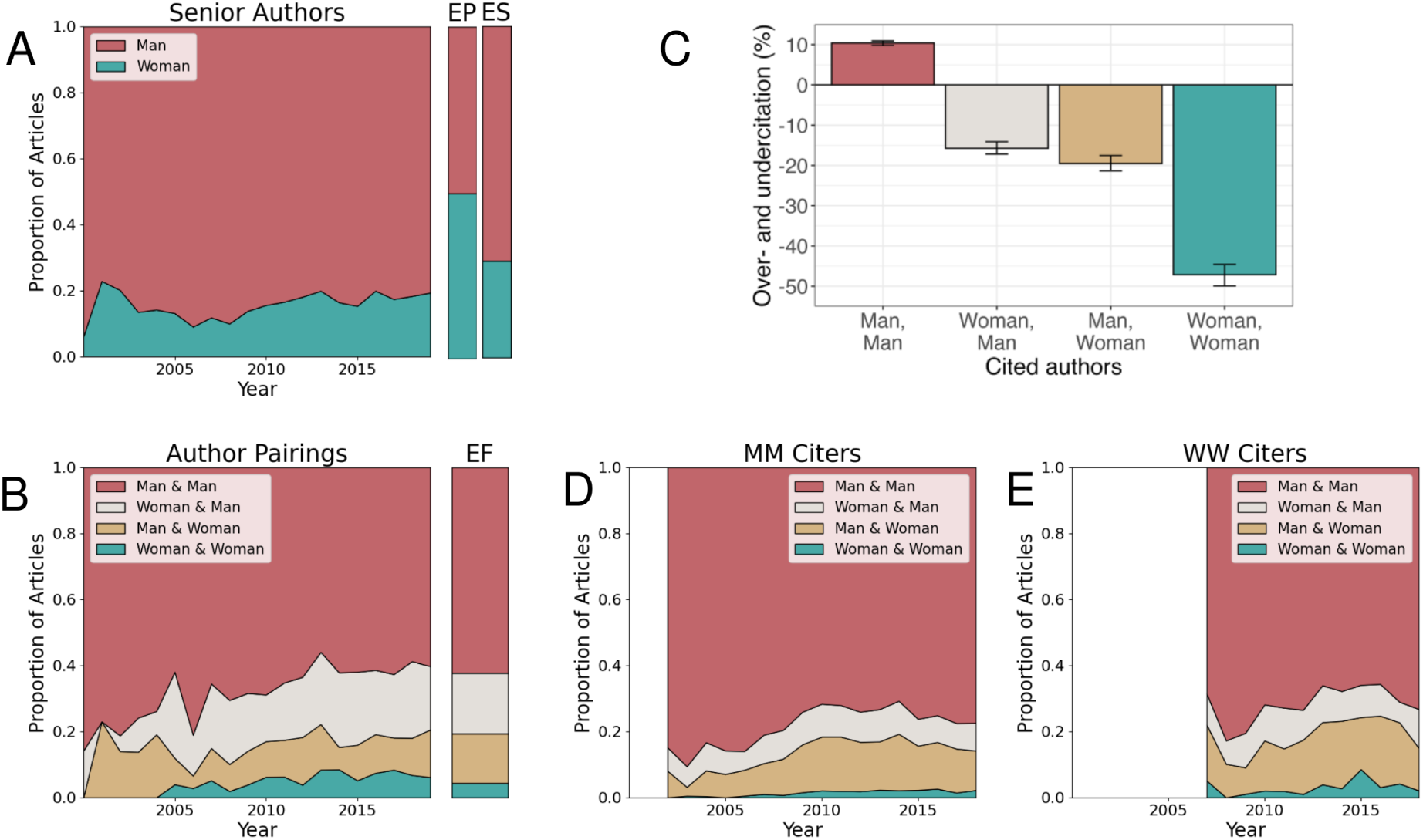
Authorship and citation results remain biased towards men authors even with a different gender inference model. [29]. (A) Proportion of articles senior-authored by each gender from 2000 to 2019. Expected values (EP) based on global proportions in 2019 and based on the Global STEM workforce (ES) in 2023 [39] are shown in the color bars at right. (B) Proportion of articles authored by a given (lead & senior author) gender pair from 2000 to 2019. Expected values (EF) produced based on the gender composition of the field in 2019 are shown in the color bar at right. (C) Rates of over- and undercitation by article lead and senior author gender based on authorship gender composition prior to publication date. (D) Authorship gender breakdown of articles cited in articles by men lead and senior authors from 2000 to 2019. (E) Authorship gender breakdown of articles cited in articles written by women lead and senior authors from 2005 to 2019.

**Figure S14.**
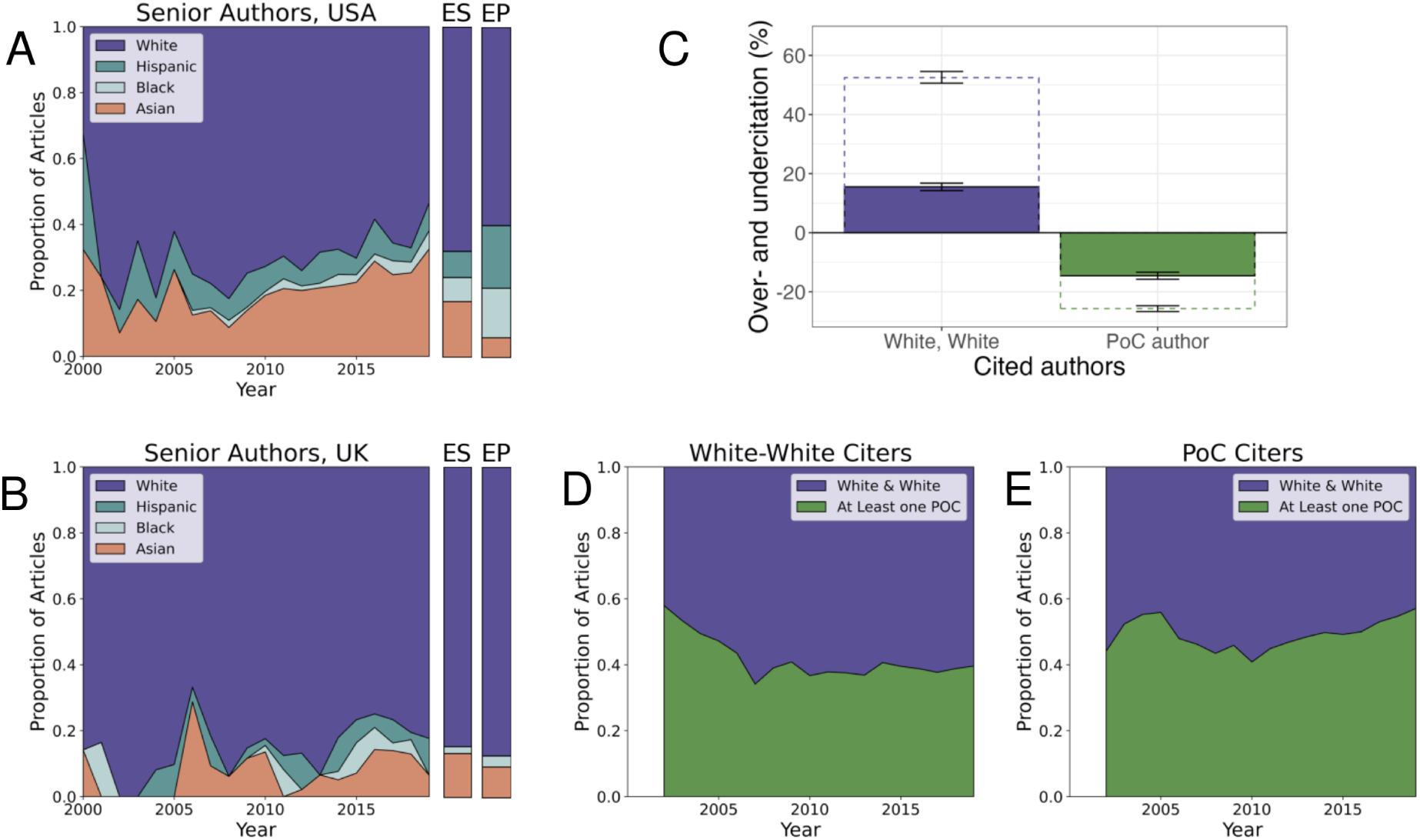
Authorship and citation practices remain biased towards White authors even with a different race/ ethnicity inference model. [29]. (A) Proportion of articles senior-authored by each race/ethnicity group in the USA from 2000 to 2019. Expected values (ES) based on the national STEM workforce based on 2019 National Science Foundation reporting [40], and expected values (EP) based on 2019 Census data [41] are shown in color bar to the right. (B) Proportion of articles senior-authored by each race/ethnicity group in the UK from 2000 to 2019. Expected values (ES) based on the national STEM academic staff [42] in 2019 and expected values (EP) based on 2019 Office of National Statistics Data [43] are shown in the color bars to the right. (C) Rates of over- and undercitation by article lead and senior author race/ethnicity based on authorship race/ethnicity composition prior to publication date. PoC author denotes that the lead and/or senior author was a person of color; sample size was too small to disaggregate by the author of color’s position. Filled boxes denote results for articles published by authors with affiliations within the Global North citing other work from the Global North. Dotted boxes denote results for articles published by authors with affiliations within the Global North citing articles with any affiliation location. (D) Racial/ethnic composition of articles cited in articles written by White lead and senior authors in the Global North, citing other articles authored in the Global North from 2002 to 2019. (E) Racial/ethnic composition of articles cited in articles written by a lead and/or senior author of color in the Global North, citing other articles authored in the Global North from 2002 to 2019.

## Appendix G: Results based on different gender/race threshold probabilities

**Figure S15.**
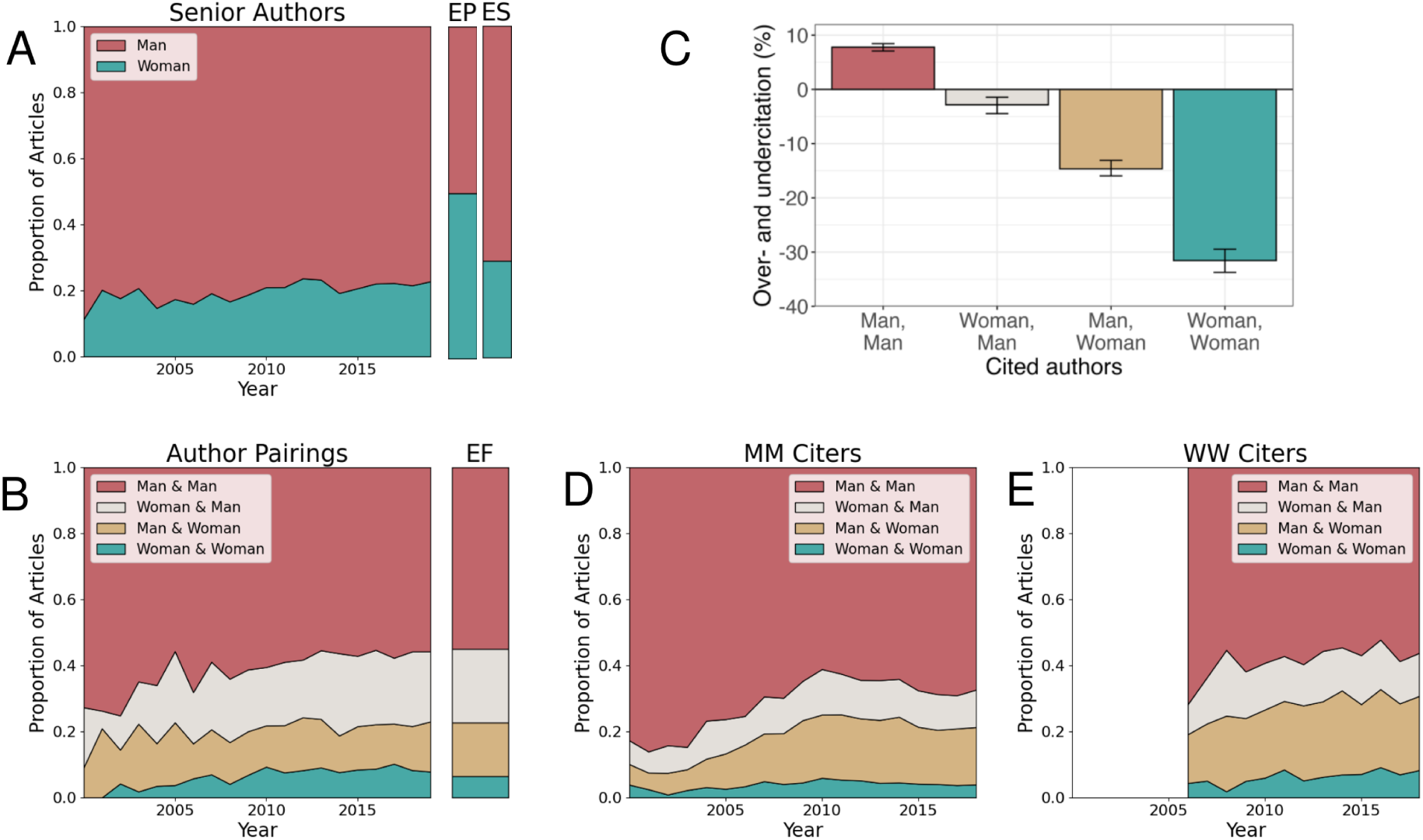
Authorship and citation practices remained biased towards men authors even with a lower, 60% probability threshold, for the gender inference model. (A) Proportion of articles senior-authored by each gender from 2000 to 2019. Expected values (EP) based on global proportions in 2019 and based on the Global STEM workforce (ES) in 2023 [39] are shown in the color bars at right. (B) Proportion of articles authored by a given (lead & senior author) gender pair from 2000 to 2019. Expected values (EF) produced based on the gender composition of the field in 2019 are shown in the color bar at right. (C) Rates of over- and undercitation by article lead and senior author gender based on authorship gender composition prior to publication date. (D) Authorship gender breakdown of articles cited in articles by men lead and senior authors from 2000 to 2019. (E) Authorship gender breakdown of articles cited in articles written by women lead and senior authors from 2006 to 2019.

**Figure S16.**
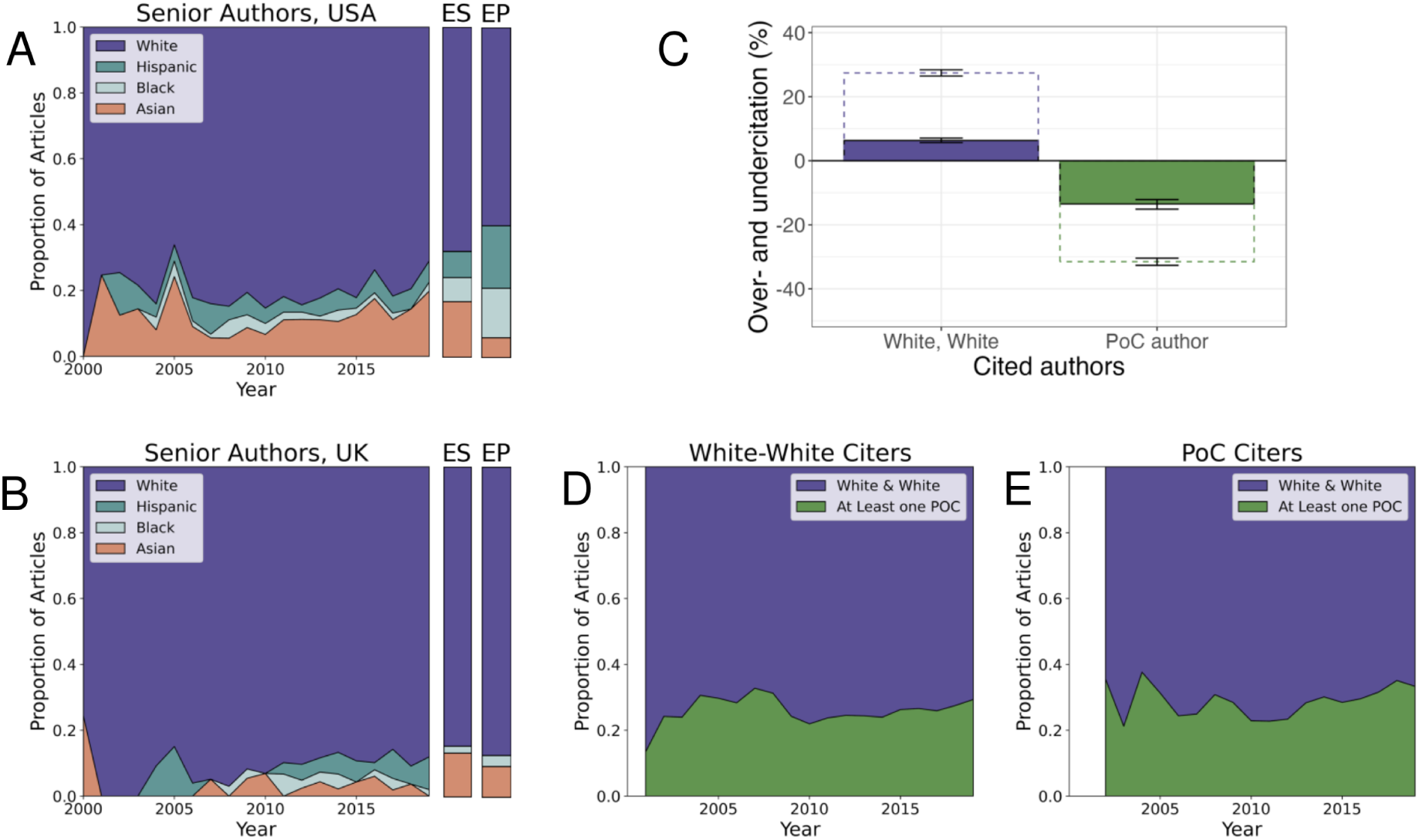
Authorship and citation practices remained biased towards White authors even with a lower, 60% probability threshold, for the race/ethnicity inference model. (A) Proportion of articles senior-authored by each race/ethnicity group in the USA from 2000 to 2019. Expected values (ES) based on the national STEM workforce based on 2019 National Science Foundation reporting [40], and expected values (EP) based on 2019 Census data [41] are shown in the color bars to the right. (B) Proportion of articles senior-authored by each race/ethnicity group in the UK from 2000 to 2019. Expected values (ES) based on the national STEM academic staff [42] in 2019, and expected values (EP) based on 2019 Office of National Statistics Data [43] are shown in the color bars to the right. (C) Rates of over- and undercitation by article lead and senior author race/ethnicity based on authorship race/ethnicity composition prior to publication date. PoC author denotes that the lead and/or senior author was a person of color; sample size was too small to disaggregate by the author of color’s position. Filled boxes denote results for articles published by authors with affiliations within the Global North citing other work from the Global North. Dotted boxes denote results for articles published by authors with affiliations within the Global North citing articles with any affiliation location. (D) Racial/ethnic composition of articles cited in articles written by White lead and senior authors in the Global North, citing other articles authored in the Global North from 2001 to 2019. (E) Racial/ethnic composition of articles cited in articles written by a lead and/or senior author of color in the Global North, citing other articles authored in the Global North from 2002 to 2019.

**Figure S17.**
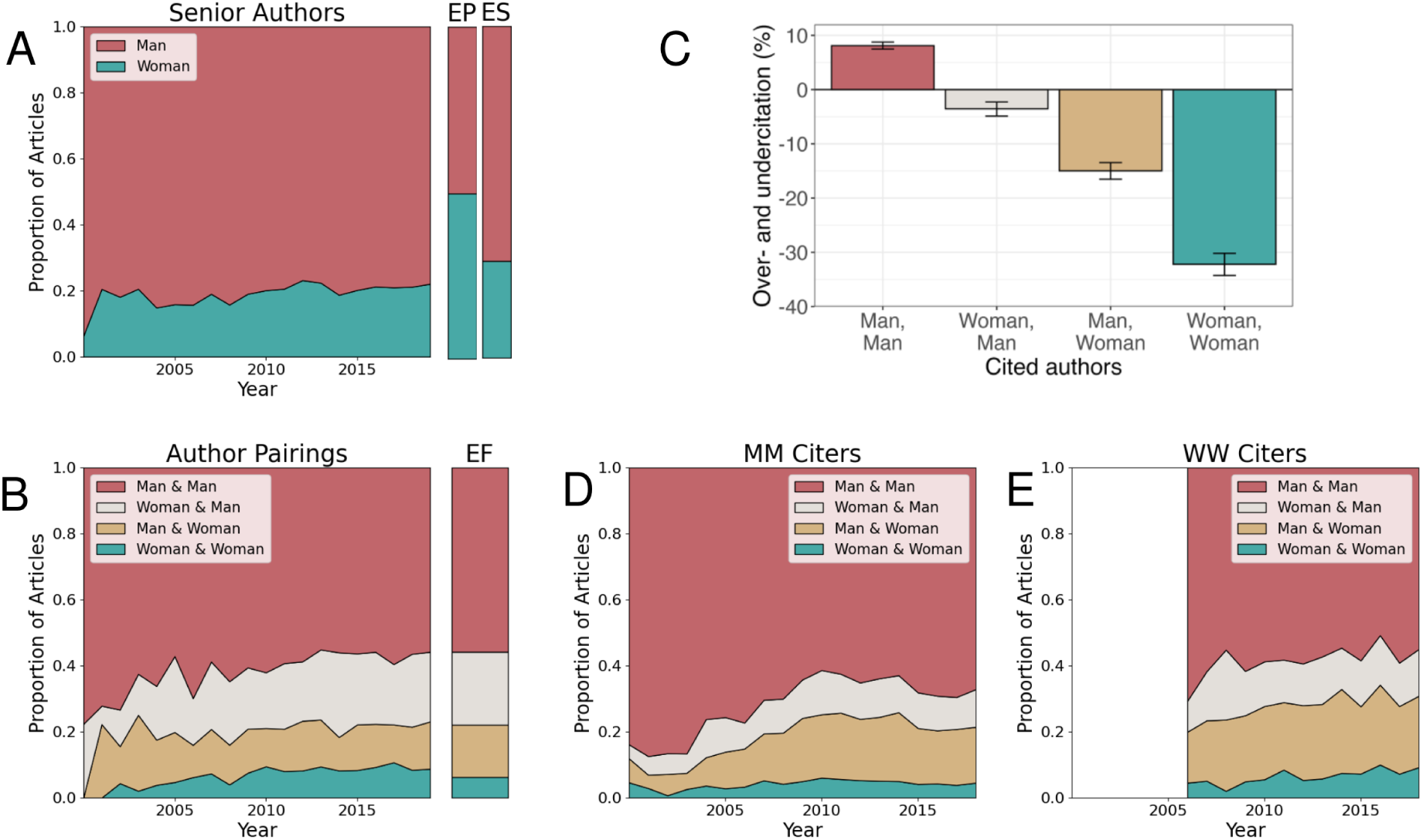
Authorship and citation practices remained biased towards men authors even with a higher, 80% probability threshold, for the gender inference model. (A) Proportion of articles senior-authored by each gender from 2000 to 2019. Expected values (EP) based on global proportions in 2019 and based on the Global STEM workforce (ES) in 2023 [39] are shown in the color bars at right. (B) Proportion of articles authored by a given (lead & senior author) gender pair from 2000 to 2019. Expected values (EF) produced based on the gender composition of the field in 2019 are shown in the color bar at right. (C) Rates of over- and undercitation by article lead and senior author gender based on authorship gender composition prior to publication date. (D) Authorship gender breakdown of articles cited in articles by men lead and senior authors from 2000 to 2019. (E) Authorship gender breakdown of articles cited in articles written by women lead and senior authors from 2006 to 2019.

**Figure S18.**
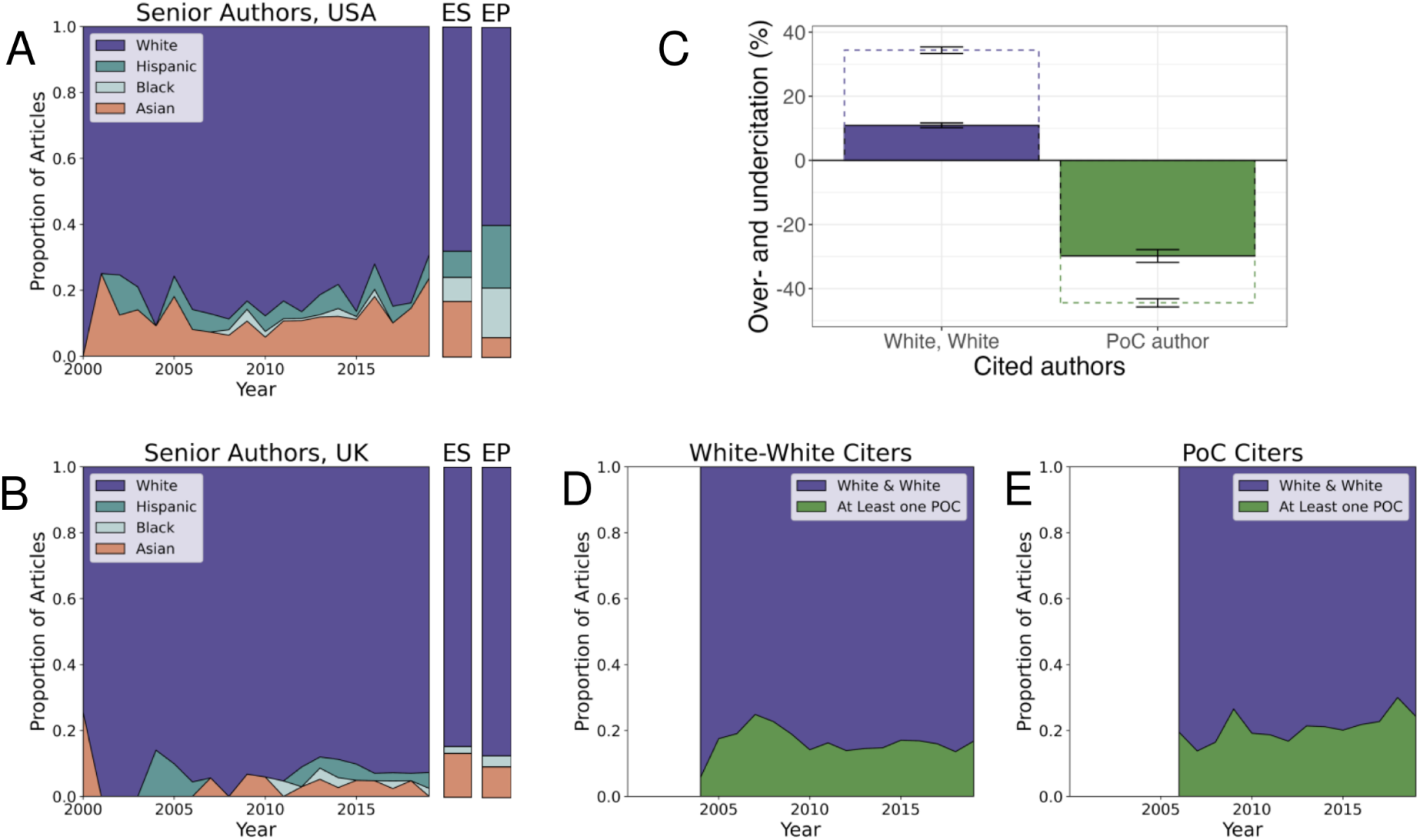
Authorship and citation practices remained biased towards White authors even with higher, 80% probability threshold, for race/ethnicity inference model. (A) Proportion of articles senior-authored by each race/ethnicity group in the USA from 2000 to 2019. Expected values (ES) based on the national STEM workforce based on 2019 National Science Foundation reporting [40], and expected values (EP) based on 2019 Census data [41] are shown in the color bars to the right. (B) Proportion of articles senior-authored by each race/ethnicity group in the UK from 2000 to 2019. Expected values (ES) based on the national STEM academic staff [42] in 2019, and expected values (EP) based on 2019 Office of National Statistics Data [43] are shown in the color bars to the right. (C) Rates of over- and undercitation by article lead and senior author race/ethnicity based on authorship race/ethnicity composition prior to publication date. PoC author denotes that the lead and/or senior author was a person of color; sample size was too small to disaggregate by the author of color’s position. Filled boxes denote results for articles published by authors with affiliations within the Global North citing other work from the Global North. Dotted boxes denote results for articles published by authors with affiliations within the Global North citing articles with any affiliation location. (D) Racial/ethnic composition of articles cited in articles written by White lead and senior authors in the Global North, citing other articles authored in the Global North from 2004 to 2019. (E) Racial/ethnic composition of articles cited in articles written by a lead and/or senior author of color in the Global North, citing other articles authored in the Global North from 2006 to 2019.

## Appendix H: Results based on lower threshold to be considered part of IDD field

**Figure S19.**
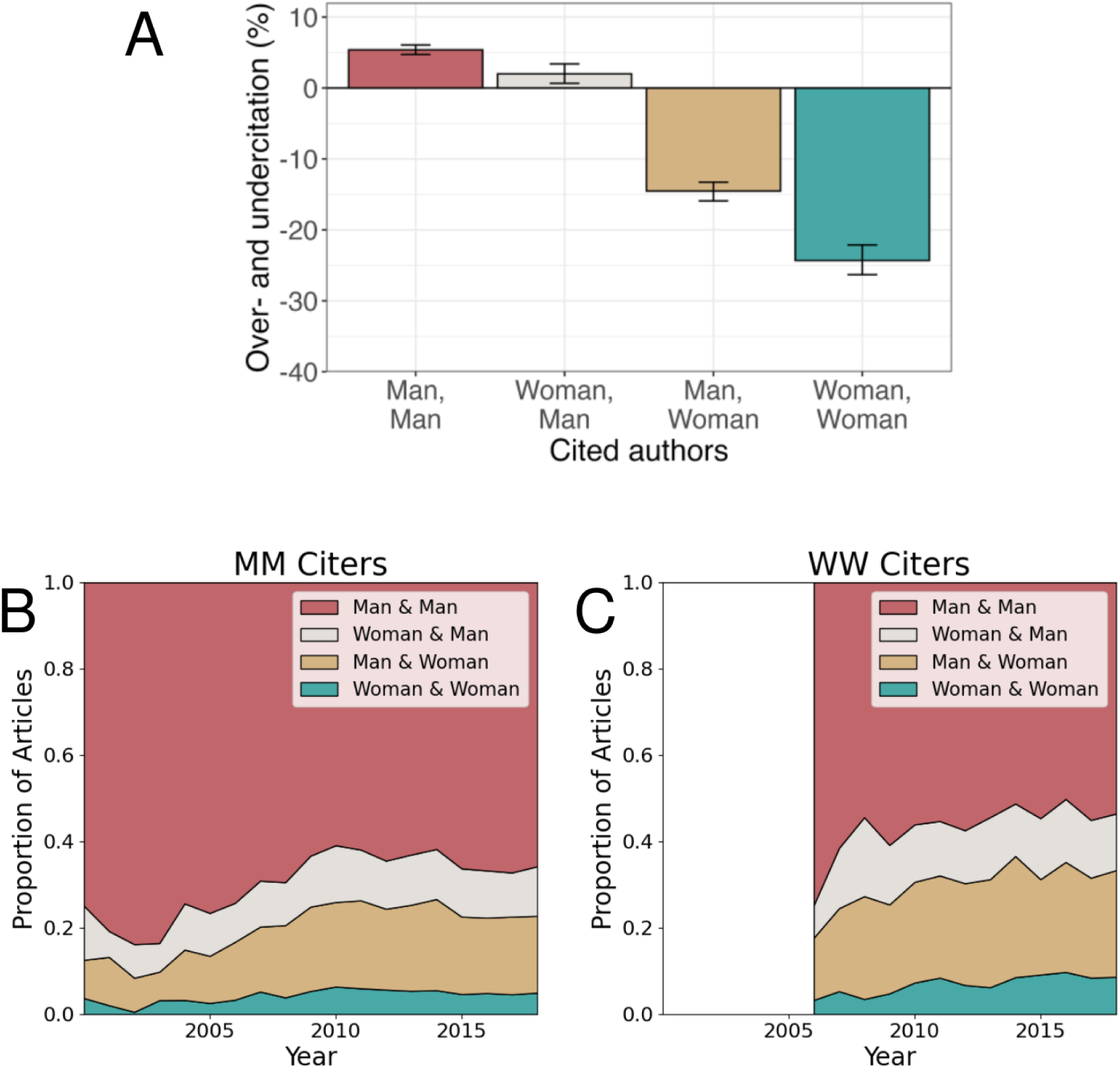
Citation practices remain biased toward articles with men lead and senior authors, despite lower, 75th percentile consensus threshold for a cited article to be considered in the field. (A) Rates of over- and undercitation by article lead and senior author gender based on authorship gender composition prior to publication date. (B) Authorship gender breakdown of articles cited in articles by men lead and senior authors from 2000 to 2019. (C) Authorship gender breakdown of articles cited in articles written by women lead and senior authors from 2006 to 2019.

**Figure S20.**
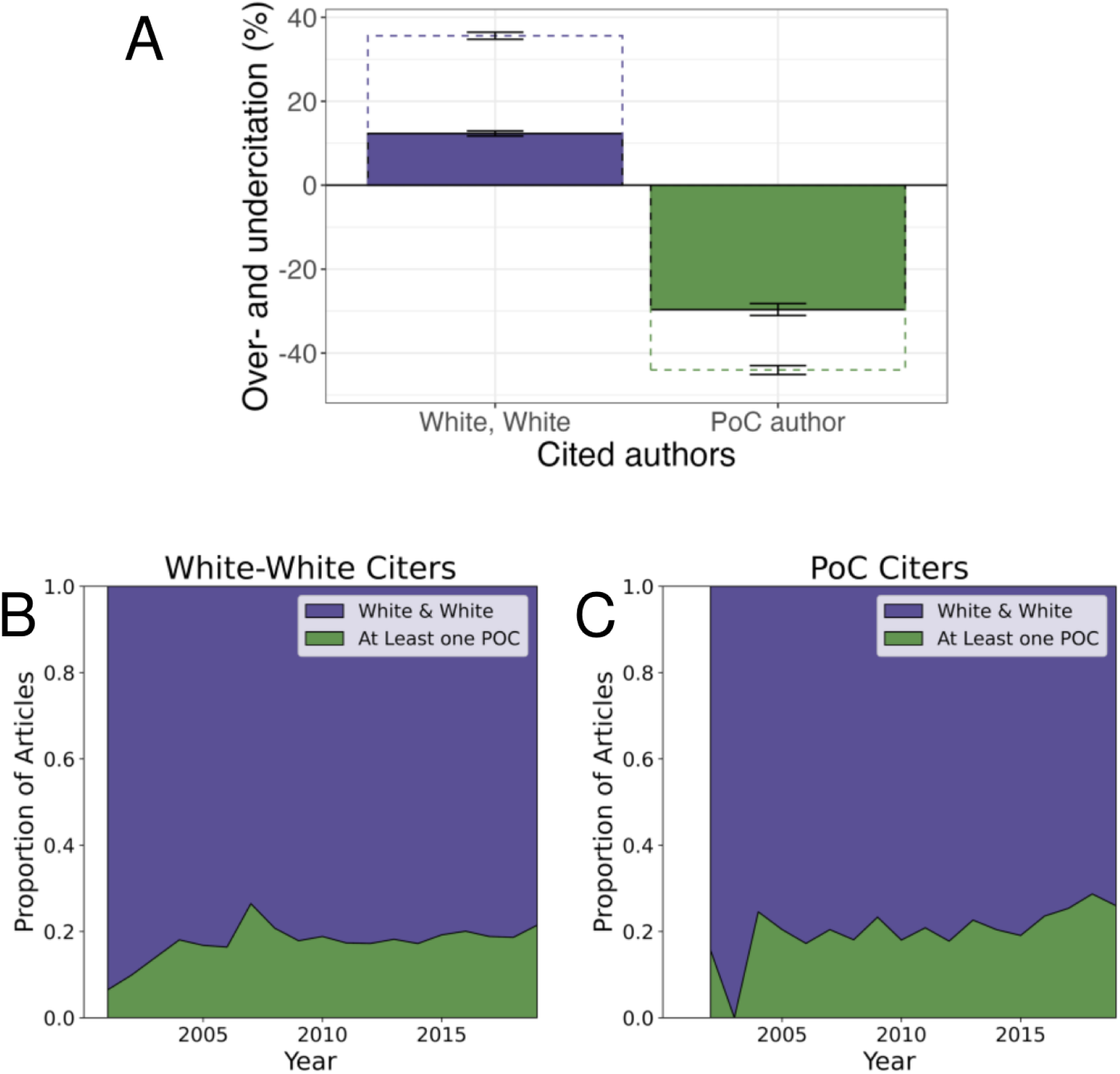
Citation practices remain biased to articles with White lead and senior authors, even with a lower, 75th percentile consensus threshold for a cited article to be considered in the field. (A) Rates of over- and undercitation by article lead and senior author race/ethnicity based on authorship race/ethnicity composition prior to publication date. PoC author denotes that the lead and/or senior author was a person of color; sample size was too small to disaggregate by the author of color’s position. Filled boxes denote results for articles published by authors with affiliations within the Global North citing other work from the Global North. Dotted boxes denote results for articles published by authors with affiliations within the Global North citing articles with any affiliation location. (B) Racial/ethnic composition of articles cited in articles written by White lead and senior authors in the Global North, citing other articles authored in the Global North from 2000 to 2019. (C) Racial/ethnic composition of articles cited in articles written by a lead and/or senior author of color in the Global North, citing other articles authored in the Global North from 2002 to 2019.

## Appendix I: Results based on the inclusion of self citations

**Figure S21.**
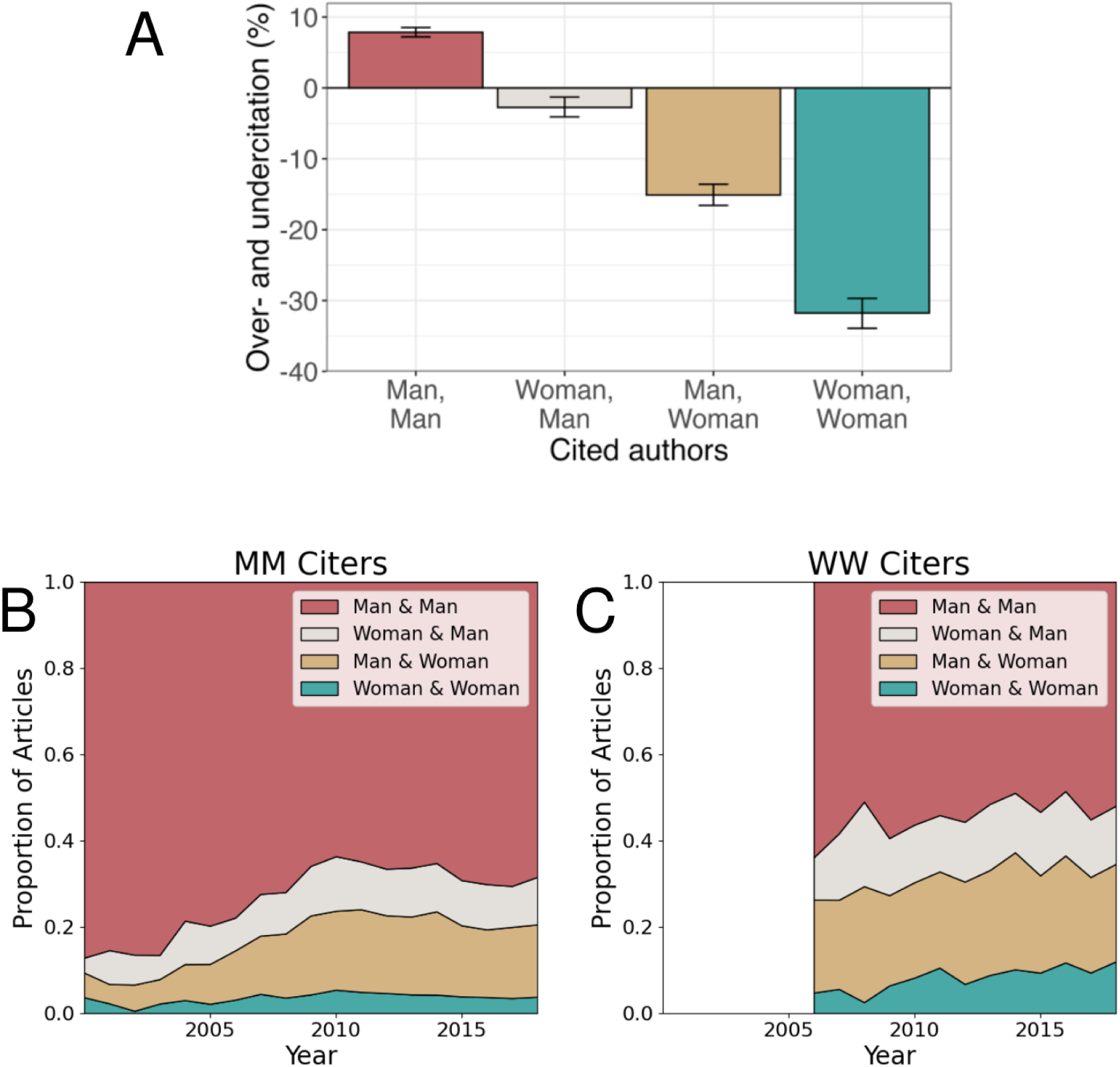
Citation practices remain biased toward articles with men lead and senior authors, with the inclusion of self citations. (A) Rates of over- and undercitation by article lead and senior author gender based on authorship gender composition prior to publication date. (B) Authorship gender breakdown of articles cited in articles by men lead and senior authors from 2000 to 2019. (C) Authorship gender breakdown of articles cited in articles written by women lead and senior authors from 2006 to 2019.

**Figure S22.**
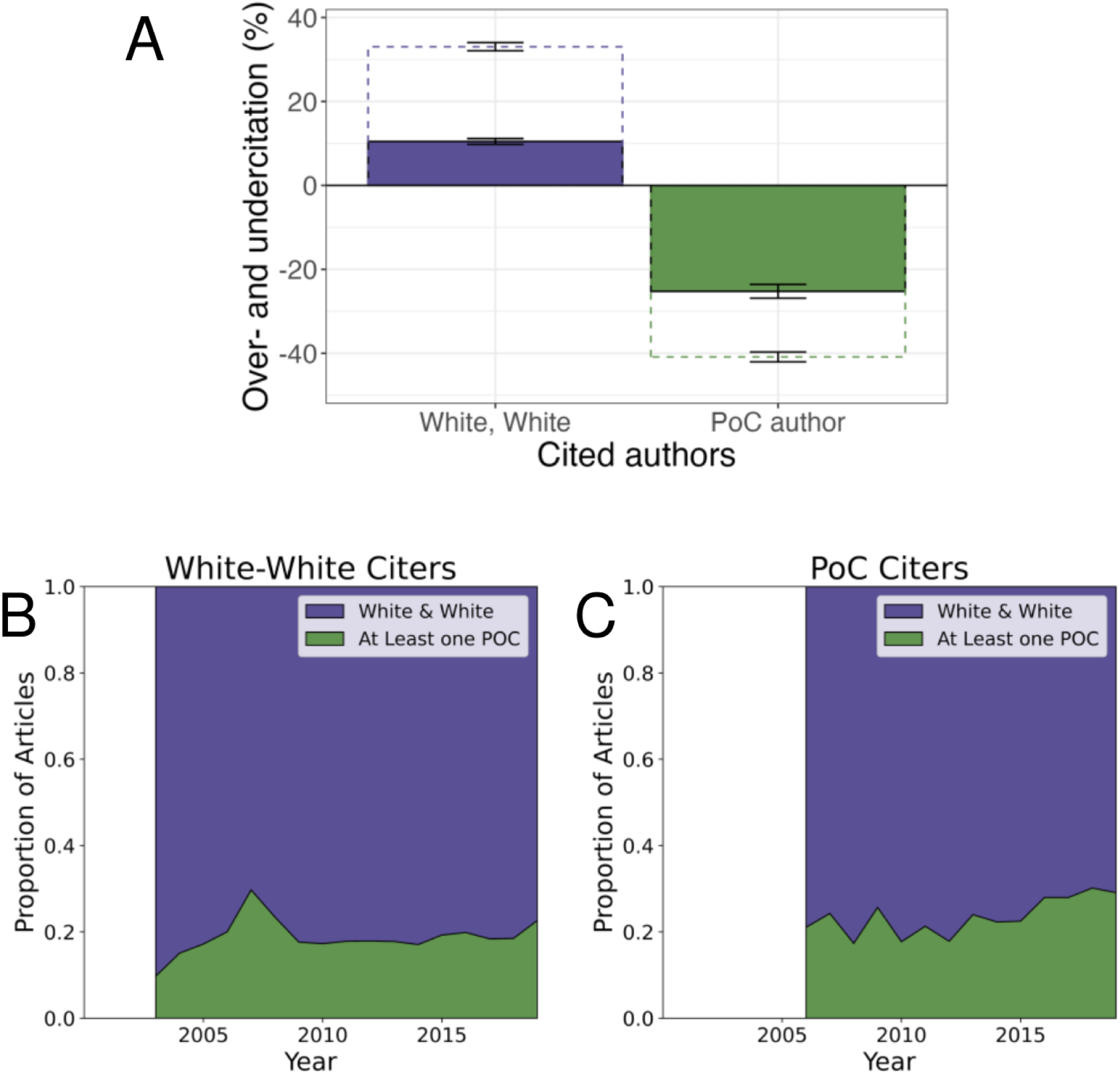
Citation practices remain biased to articles with White lead and senior authors, with the inclusion of self citations. (A) Rates of over- and undercitation by article lead and senior author race/ethnicity based on authorship race/ethnicity composition prior to publication date. PoC author denotes that the lead and/or senior author was a person of color; sample size was too small to disaggregate by the author of color’s position. Filled boxes denote results for articles published by authors with affiliations within the Global North citing other work from the Global North. Dotted boxes denote results for articles published by authors with affiliations within the Global North citing articles with any affiliation location. (B) Racial/ethnic composition of articles cited in articles written by White lead and senior authors in the Global North, citing other articles authored in the Global North from 2000 to 2019.(C) Racial/ethnic composition of articles cited in articles written by a lead and/or senior author in the Global North, citing other articles authored in the Global North of color from 2002 to 2019.

## Appendix J: Regression diagnostics and sensitivity analyses

Using a chi-squared test, we determined that neither lead and senior author gender (*χ*^2^ = 84.8, *p <* 0.0001) nor lead and senior author race are independent (*χ*^2^ = 1122.8, *p <* 0.0001) in the citation dataset. This result justified the use of author pairings in our regression analysis. By regressing square-root transformed impact factor on paired author identity, we determined that both gender and JIF (Figure S23) and race and JIF are collinear (Figure S24). These findings justify the inclusion of a interactive effect between gender and JIF, and race and JIF but warrant caution in interpreting our results. By regressing log_10_citation rate on gender alone, and race alone, we found there was a weak association between author gender and article citation rate (Figure S25) and a strong association between author race and article citation rate, if both authors were non-White (Figure S26). Impact factor could be complicating these results and, when included, suggests that articles with both lead and senior women authors (Figure S27) or authors of color (Figure S28) have lower citation rates.

These results are not sensitive to the inclusion of self-citations (Figure S30). When the analysis is restricted to articles published in the Global North or authorship articles that aren’t sufficiently cited on their own are excluded, race has a lesser or no effect on citation rate (Figure S31, S32).

**Figure S23.**
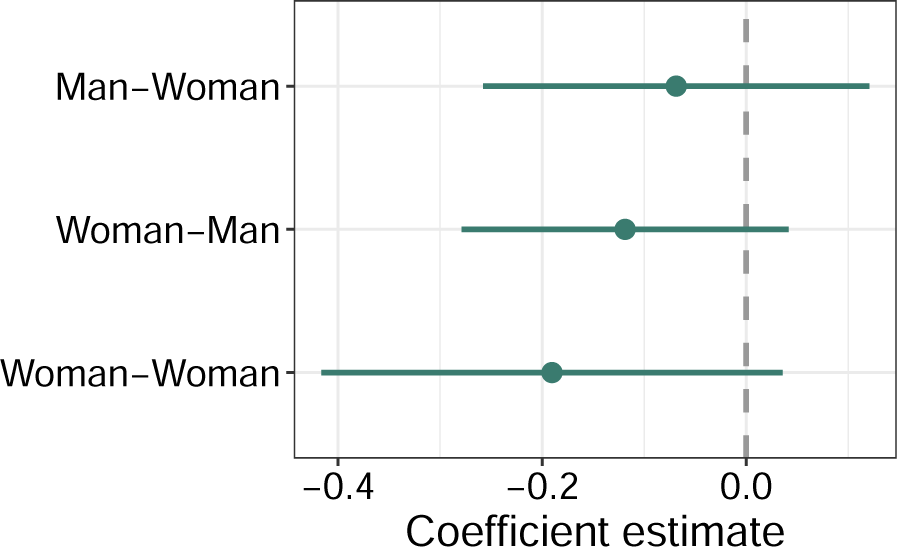
Regression of square-root transformed Journal Impact Factor on author pairing gender suggests these predictors are slightly collinear.

**Figure S24.**
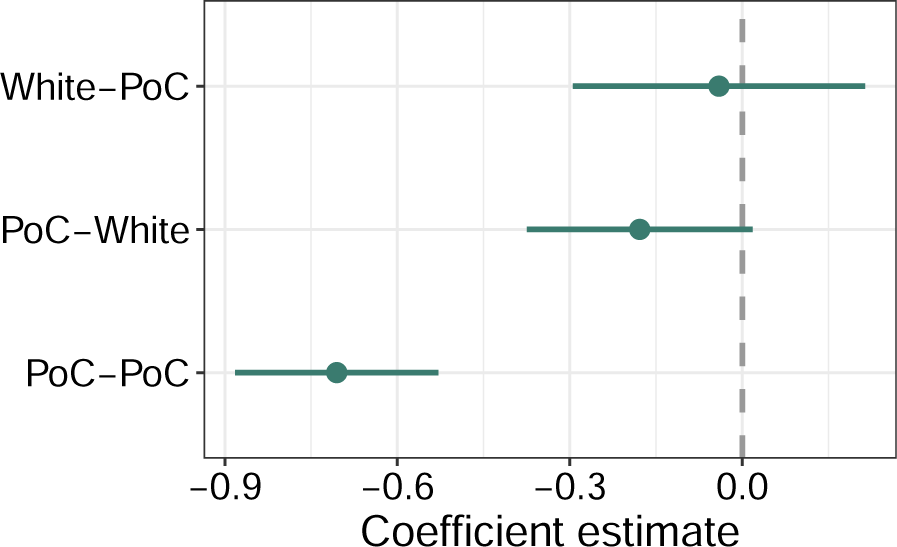
Regression of square-root transformed Journal Impact Factor on author pairing race suggests these predictors are collinear.

**Figure S25.**
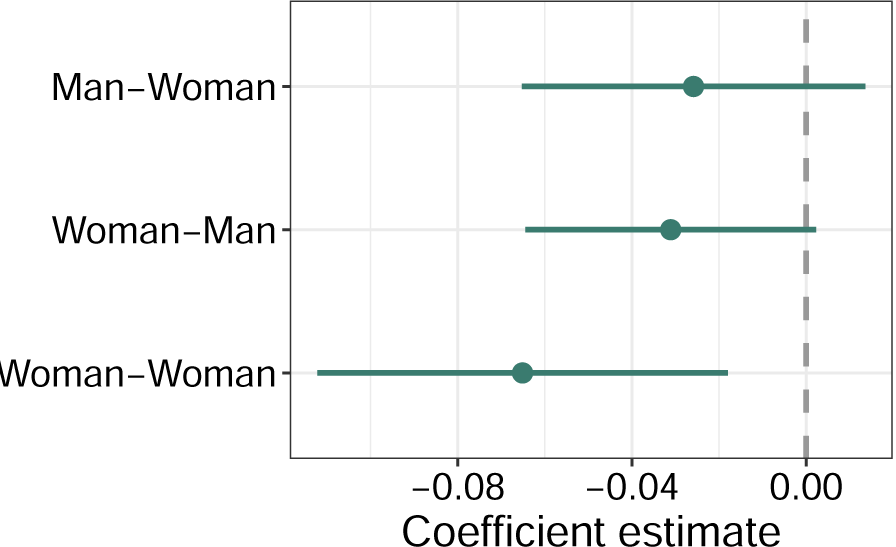
Coefficients from linear regression predicting citation rate (num. citations/years since publication) based on article and author information. This analysis excludes 2023 Journal Impact Factor.

**Figure S26.**
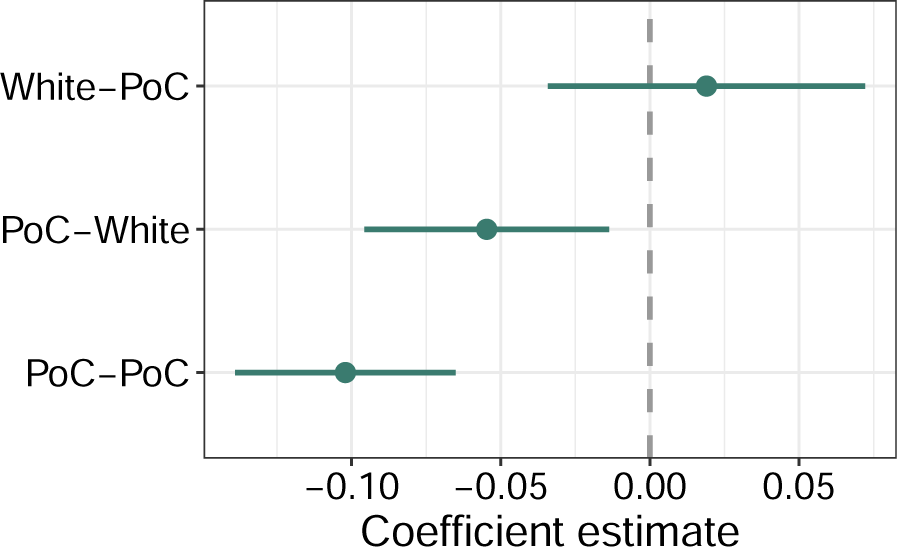
Coefficients from linear regression predicting citation rate (num. citations/years since publication) based on article and author information. PoC includes Asian, Hispanic, and Black authors. This analysis excludes 2023 Journal Impact Factor.

**Figure S27.**
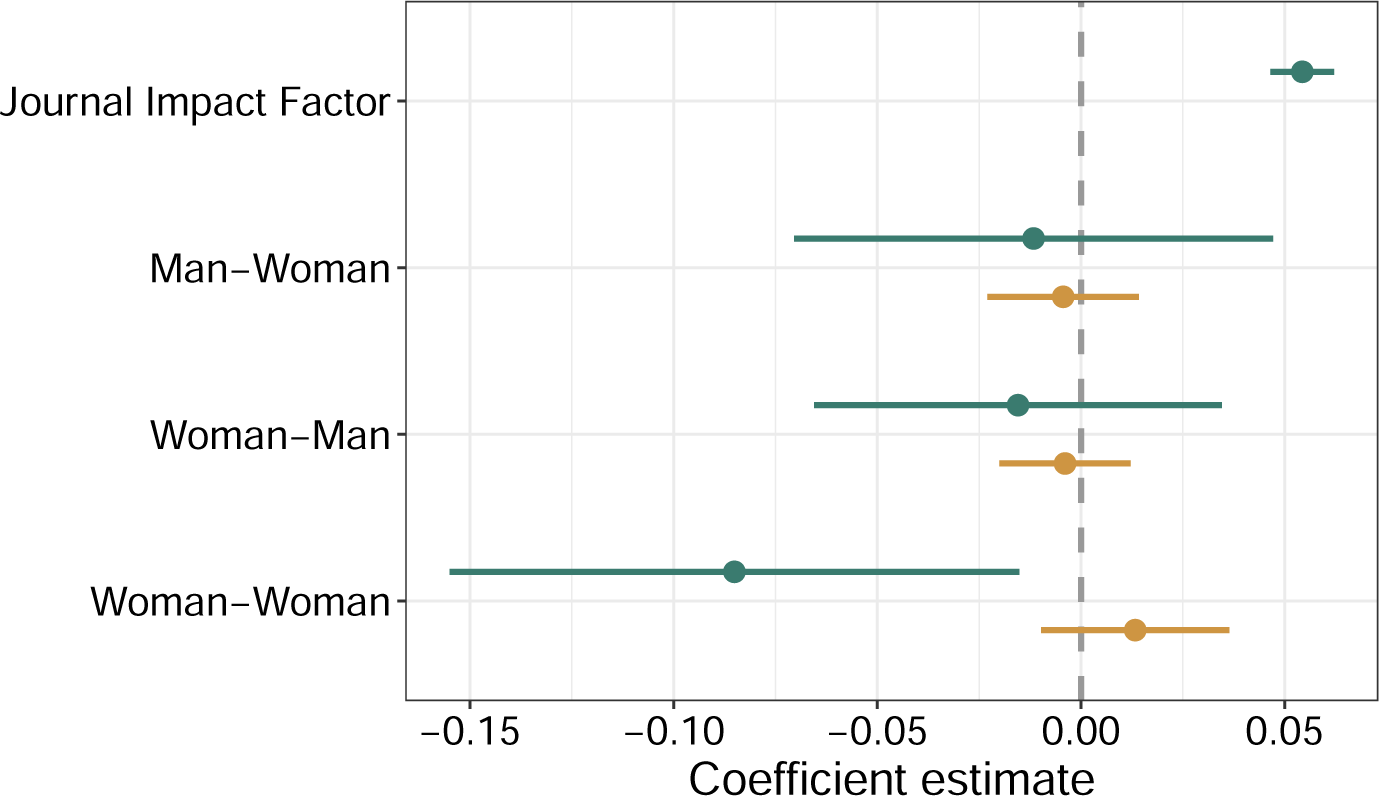
Coefficients from linear regression predicting citation rate (num. citations/years since publication) based on article and author information. This analysis ignores author race. Teal point-ranges show the main effects (coefficient estimate and 95% confidence interval) for the independent impact of the predictor on citation rates, and gold point-ranges show the interaction effects for the impact of the predictor on citation rates for different values of Journal Impact Factor.

**Figure S28.**
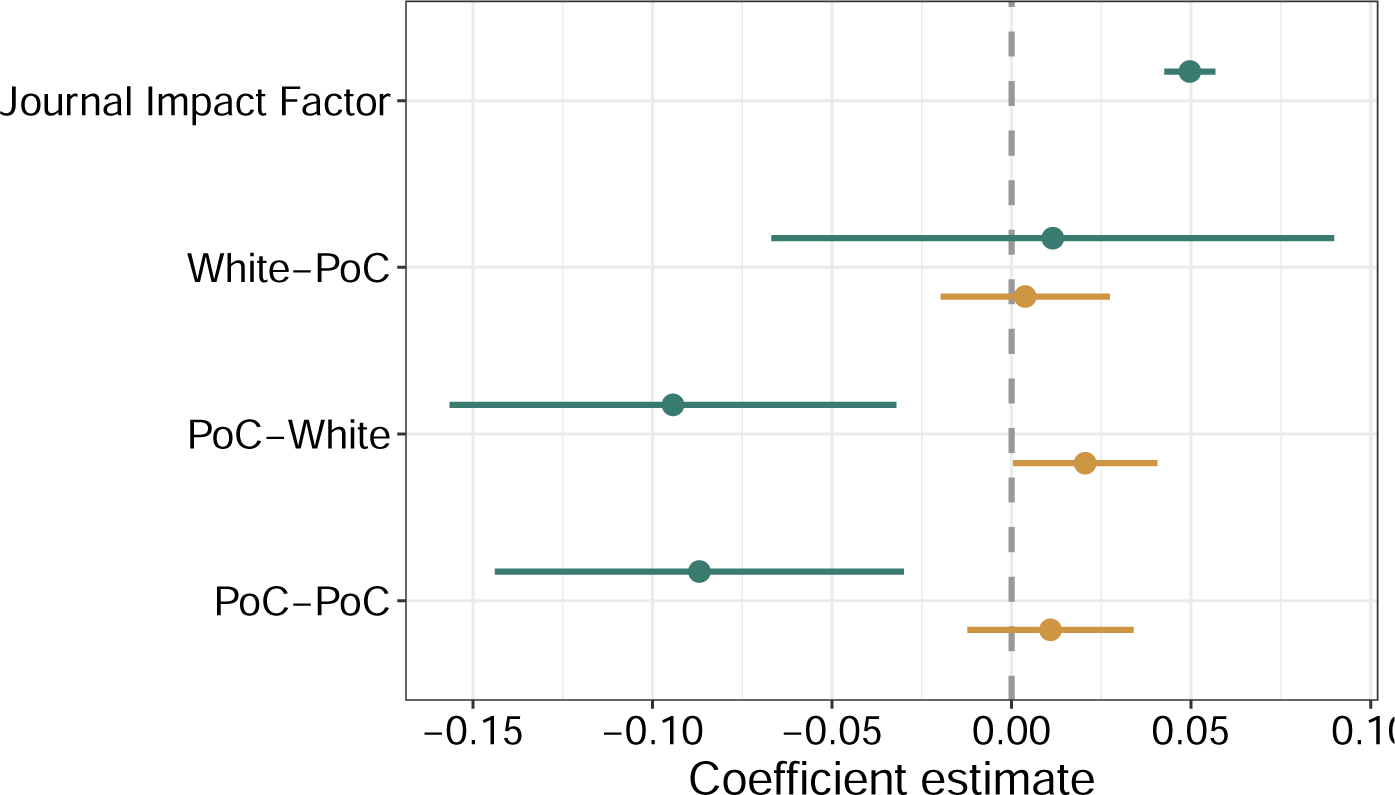
Coefficients from linear regression predicting citation rate (num. citations/years since publication) based on article and author information. PoC includes Asian, Hispanic, and Black authors. This analysis ignores author gender. Teal point-ranges show the main effects (coefficient estimate and 95% confidence interval) for the independent impact of the predictor on citation rates, and gold point-ranges show the interaction effects for the impact of the predictor on citation rates for different values of Journal Impact Factor.

**Figure S29.**
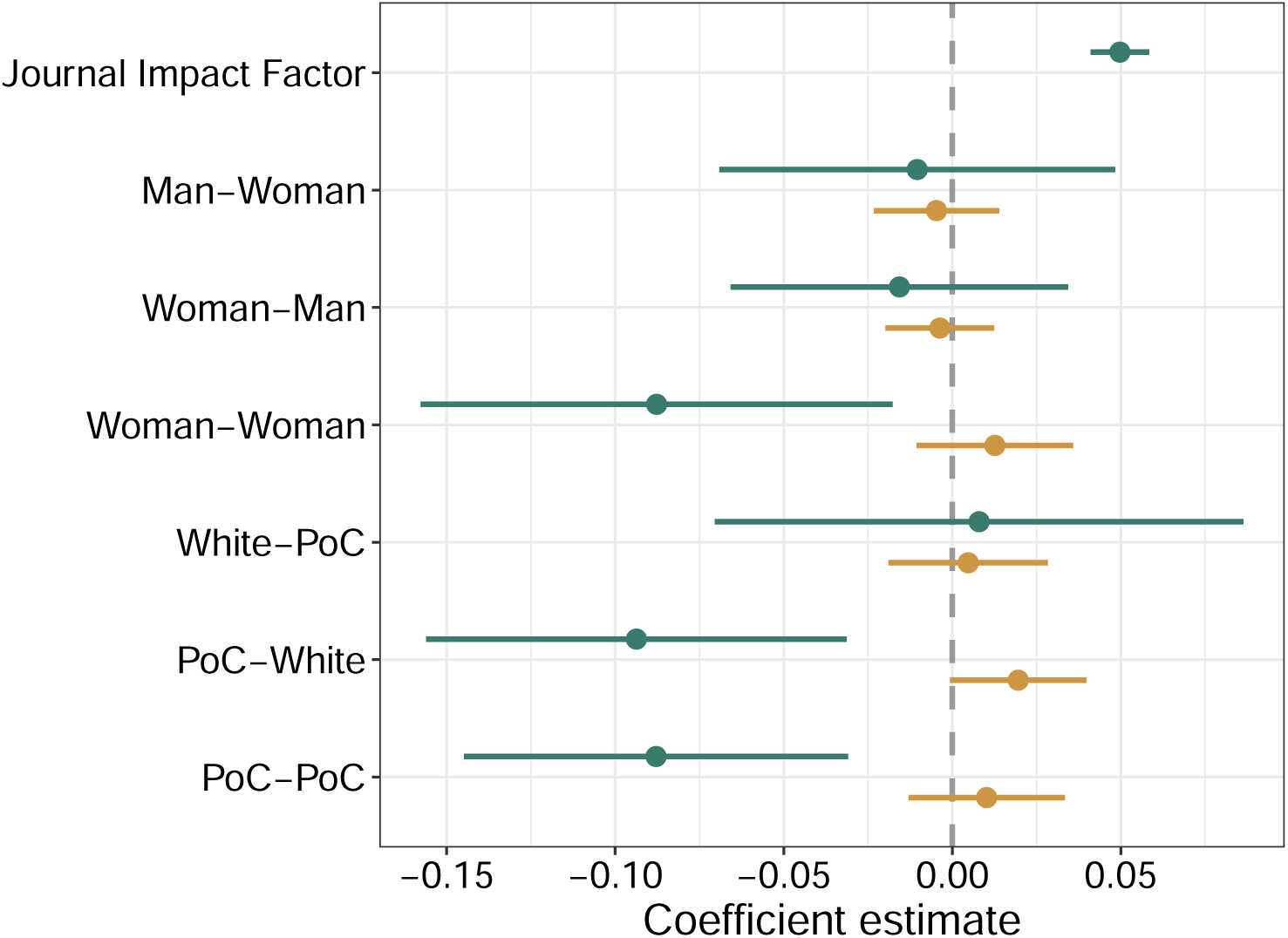
Coefficients from linear regression predicting citation rate (num. citations/years since publication) based on article and author information. PoC includes Asian, Hispanic, and Black authors. This is identical to the regression in the main text with the exclusion of a Global North indicator covariate. Teal point-ranges show the main effects (coefficient estimate and 95% confidence interval) for the independent impact of the predictor on citation rates, and gold point-ranges show the interaction effects for the impact of the predictor on citation rates for different values of Journal Impact Factor.

**Figure S30.**
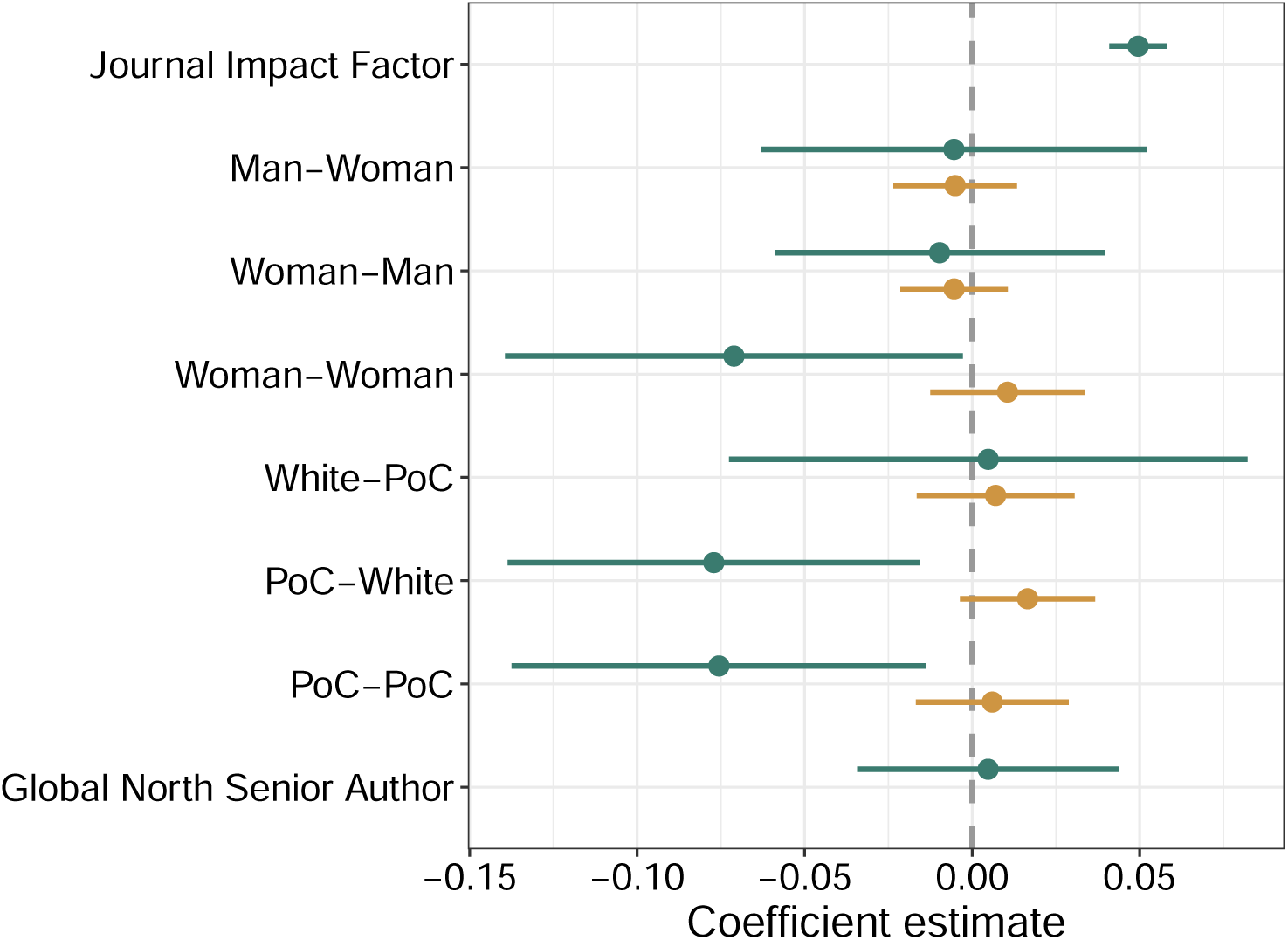
Coefficients from linear regression predicting citation rate (num. citations/years since publication) based on article and author information. PoC includes Asian, Hispanic, and Black authors. This analysis permits self-citations. Teal point-ranges show the main effects (coefficient estimate and 95% confidence interval) for the independent impact of the predictor on citation rates, and gold point-ranges show the interaction effects for the impact of the predictor on citation rates for different values of Journal Impact Factor.

**Figure S31.**
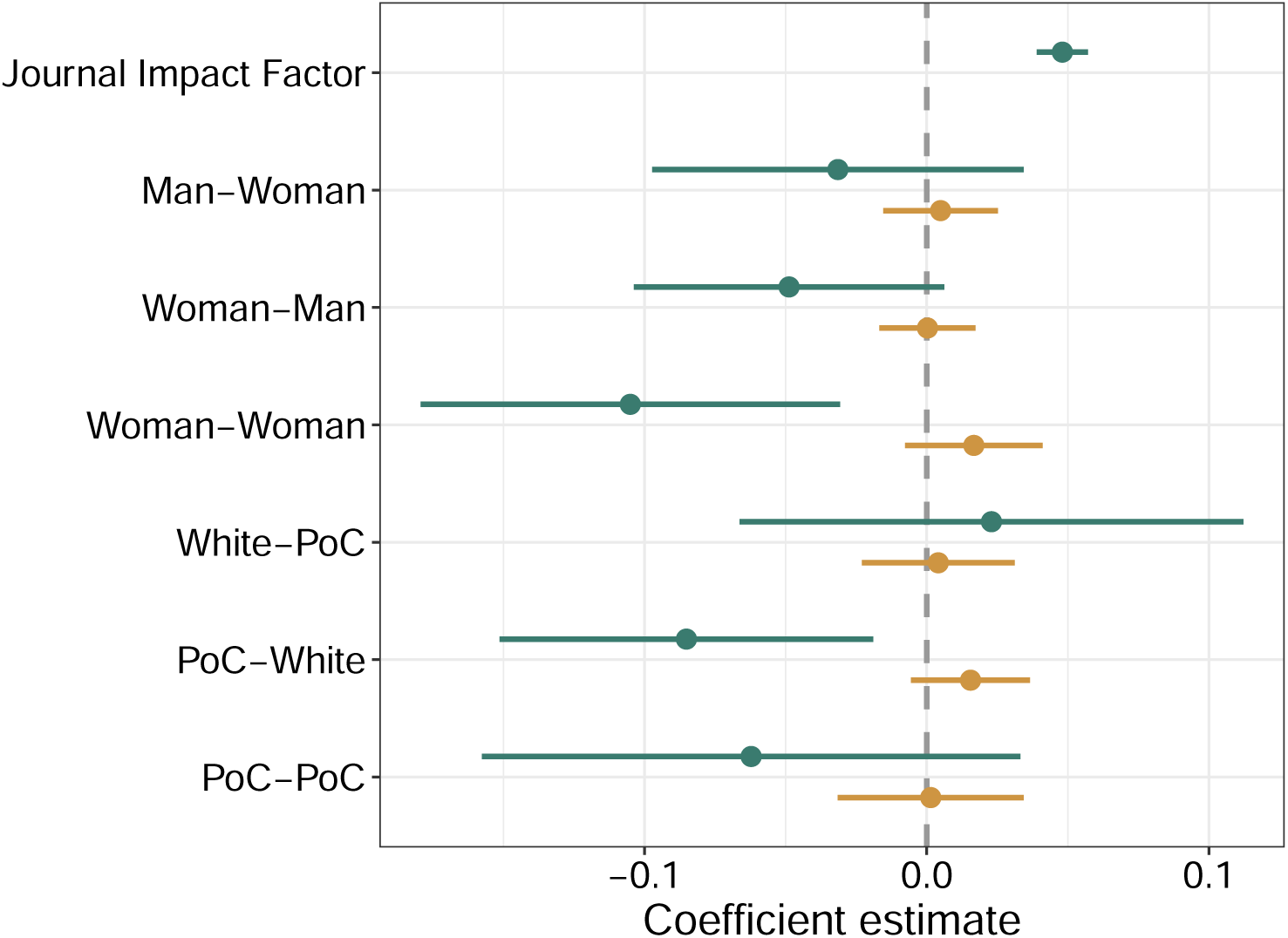
Coefficients from linear regression predicting citation rate (num. citations/years since publication) based on article and author information. PoC included Asian, Hispanic, and Black authors. Limited to articles written by senior authors in the Global North. Teal point-ranges show the main effects (coefficient estimate and 95% confidence interval) for the independent impact of the predictor on citation rates, and gold point-ranges show the interaction effects for the impact of the predictor on citation rates for different values of Journal Impact Factor.

**Figure S32.**
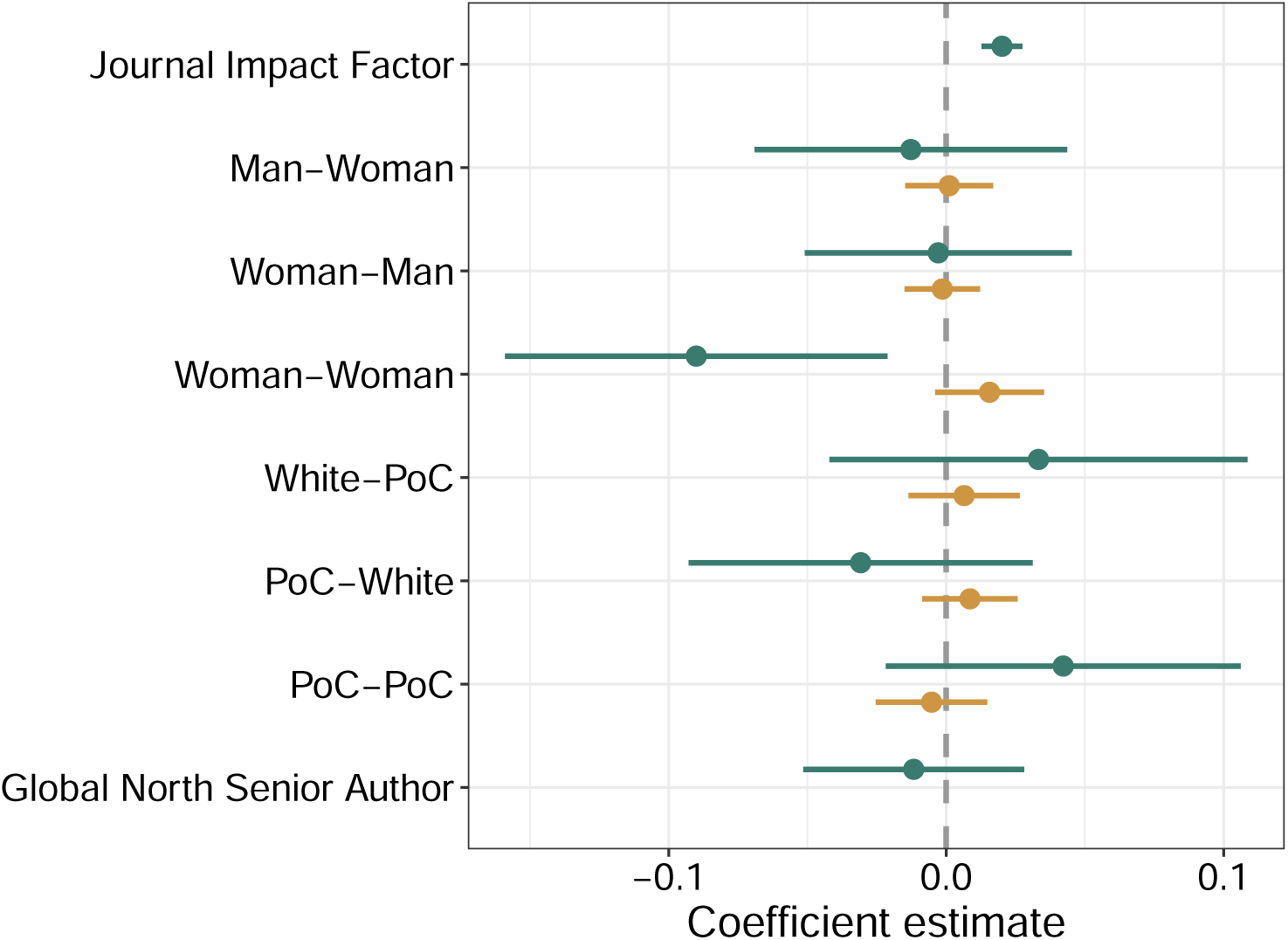
Coefficients from linear regression predicting citation rate (num. citations/years since publication) based on article and author information. PoC includes Asian, Hispanic, and Black authors. Articles must be sufficiently cited as defined in the manuscript, with no exceptions for articles in the authorship dataset that have fewer citations. Teal point-ranges show the main effects (coefficient estimate and 95% confidence interval) for the independent impact of the predictor on citation rates, and gold point-ranges show the interaction effects for the impact of the predictor on citation rates for different values of Journal Impact Factor.

## Appendix K: Intersectional results

**Figure S33.**
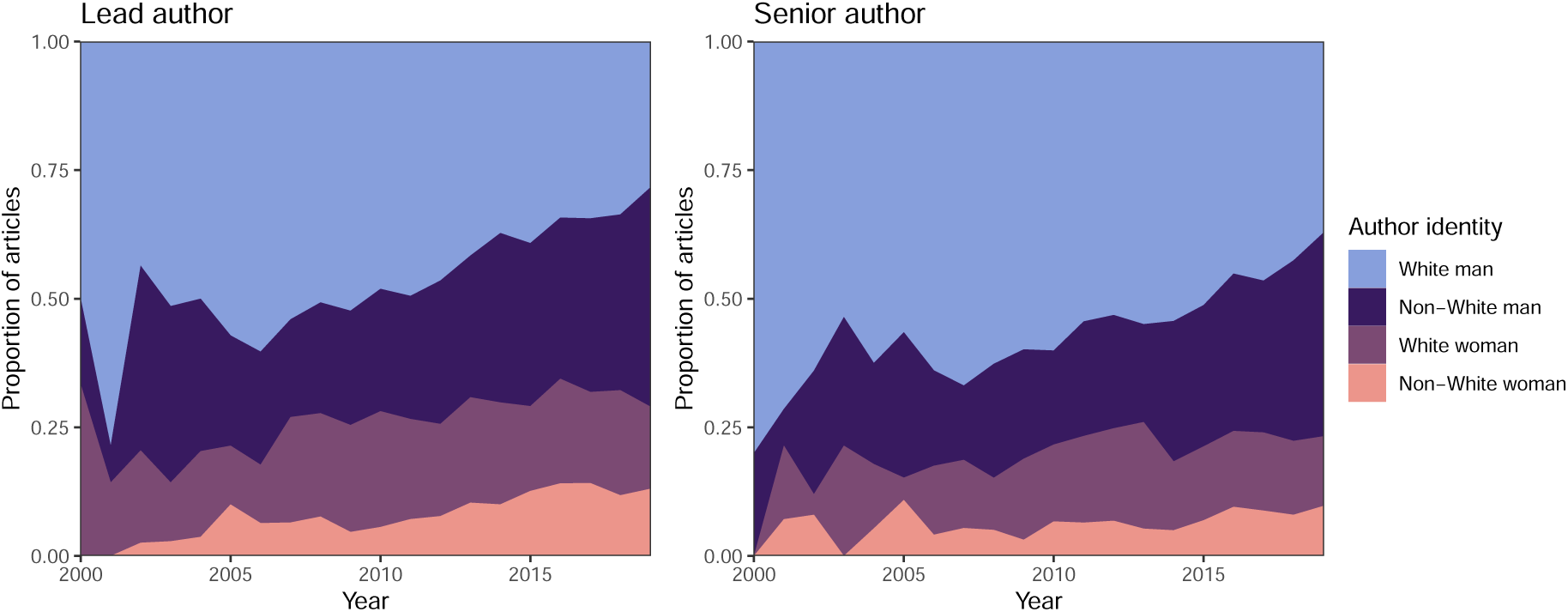
Intersectional authorship composition for lead and senior authors over time. This analysis uses the global authorship dataset from the main text.

